# Stimulating the *sir2-pgc-1ɑ* axis rescues exercise capacity and mitochondrial respiration in *Drosophila tafazzin* mutants

**DOI:** 10.1101/2022.01.06.475267

**Authors:** Deena Damschroder, Rubén Zapata-Pérez, Riekelt H. Houtkooper, Robert Wessells

## Abstract

Cardiolipin (CL) is a phospholipid required for proper mitochondrial function. Tafazzin remodels CL to create highly unsaturated fatty acid chains. However, when *tafazzin* is mutated, CL remodeling is impeded, leading to mitochondrial dysfunction and the disease Barth syndrome. Patients with Barth syndrome often have severe exercise intolerance, which negatively impacts their overall quality of life. Boosting NAD^+^ levels can improve symptoms of other mitochondrial diseases, but its effect in the context of Barth syndrome has not been examined. We demonstrate for the first time that nicotinamide riboside (NR) can rescue exercise tolerance and mitochondrial respiration in a *Drosophila tafazzin* mutant and that the beneficial effects are dependent on *sir2* and *pgc-1α*. Overexpressing *pgc-1α* increased the total abundance of cardiolipin in mutants. In addition, muscles and neurons were identified as key targets for future therapies because *sir2* or *pgc-1α* overexpression in either of these tissues is sufficient to restore the exercise capacity of *Drosophila tafazzin* mutants.

**Summary Statement:** Nicotinamide riboside rescues the exercise capacity and mitochondrial function of a *Drosophila* model of Barth syndrome in a Sir2/Pgc-1ɑ dependent manner.

## Introduction

The mitochondrial-specific phospholipid cardiolipin (CL) has a substantial impact on mitochondrial metabolism. CL supports mitochondrial function in multiple ways including directly binding to the complexes of the respiratory chain (Fry and Green, 1980, Schlame and Haldar, 1993, Pfeiffer et al., 2003) and assisting with the formation of super complexes to allow efficient oxidative phosphorylation (Pfeiffer et al., 2003, Mckenzie et al., 2006). CL is synthesized within the inner mitochondrial membrane and *de novo* CL undergoes a remodeling process that alters the CL lipid profile to contain mostly unsaturated fatty acids (Hostetler et al., 1971, Schlame et al., 2005, Schlame, 2013).

Tafazzin is the primary enzyme that remodels CL and loss of function mutations in *tafazzin* increases the ratio of monolysocardiolipin (MLCL) to CL (Vreken et al., 2000). This alteration in CL leads to mitochondrial dysfunction and a rare disease known as Barth syndrome (Barth et al., 1983, Vreken et al., 2000, Schlame et al., 2002). Barth syndrome is an X-linked disorder that is characterized by cardiomyopathy, neutropenia, muscle weakness, delayed growth, increased urinary 3-methylglutaconic aciduria, and exercise intolerance (Neustein et al., 1979, Barth et al., 1983, Kelley et al., 1991, Roberts et al., 2012, Cade et al., 2017, Bittel et al., 2018), with patients reporting that their exercise intolerance is one of the principal symptoms that most negatively impact their lives (Bowen et al., 2019).

Barth syndrome currently has no cure and patient care focuses on symptom management, making continued efforts to find treatments to reduce the severity of symptoms imperative. One possible therapeutic avenue is manipulating the levels of nicotinamide adenine dinucleotide (NAD^+^). A decreased NAD^+^ concentration is associated with many mitochondrial diseases, age-related diseases, and general aging (Massudi et al., 2012, Clement et al., 2019, Fang et al., 2019, Pirinen et al., 2020, Zapata-Perez et al., 2021). In various disease states, supplementation with NAD^+^ precursors such as nicotinamide riboside (NR), nicotinic acid (NA), and nicotinamide mononucleotide (NMN) provides improvements to mitochondrial function in flies (Lehmann et al., 2017) and in mice (Khan et al., 2014, Zhou et al., 2020), highlighting the importance of maintaining NAD^+^ homeostasis.

There are no major side effects reported with NR supplementation (Martens et al., 2018, Conze et al., 2019), although supplementation with NA can cause uncomfortable skin flushing (Kamanna et al., 2009). Both NMN and NR supplementation can increase NAD^+^ levels, but the oral bioavailability and transport into cells differs between the two compounds, with NR transported directly into the cell, while NMN is either converted to NR by CD73 and then transported into the cell (Garavaglia et al., 2012, Grozio et al., 2013, Ratajczak et al., 2016, Rajman et al., 2018) or directly transported by Slc12a8 in the small intestine (Grozio et al., 2019). Chronic NR and NMN supplementation is well tolerated in humans (Conze et al., 2016, Trammell et al., 2016, Martens et al., 2018, Conze et al., 2019, Elhassan et al., 2019, Remie et al., 2020, Yoshino et al., 2021).

Sirtuins are known for their role in longevity and mitochondrial health (Li et al., 2008, Fang et al., 2016, Imai and Guarente, 2016, Lautrup et al., 2019, Zapata-Perez et al., 2021). Dietary supplementation with NMN or NR can increase the activity of Sir2 (Yoshino et al., 2011, Canto et al., 2012, Wang et al., 2018). Among its many targets, Sir2 can deacetylate Pgc-1α and increase its activity (Rodgers et al., 2005, Imai and Guarente, 2016). Pgc-1α is a well-studied transcription coactivator that regulates the expression of mitochondrial oxidative phosphorylation genes to improve mitochondrial function and increase mitochondrial biogenesis (Wu et al., 1999, Lehman et al., 2000, Lai et al., 2014, Imai and Guarente, 2016). PGC-1ɑ is upregulated with exercise training and upregulates antioxidant defense systems to limit oxidative damage from increased mitochondrial respiration (Handschin et al., 2007, Austin and St-Pierre, 2012).

The function of the *Drosophila pgc-1α* homolog, *spargel*, is similar to its mammalian counterpart, including its role in longevity, mitochondrial biogenesis, oxidative stress resistance, and exercise adaptations (Tiefenbock et al., 2010, Rera et al., 2011, Tinkerhess et al., 2012b, Mukherjee et al., 2014). In the context of Barth syndrome, overexpressing either *sir2* or *pgc-1α* in a *Drosophila tafazzin* mutant can alter the cardiolipin profile to more wild-type levels (Xu et al., 2019). However, the impact of these changes on exercise tolerance and mitochondrial respiration has not previously been examined.

We demonstrate for the first time the beneficial effects of NR administration in the context of Barth syndrome. Supplementation with NR was sufficient to restore the endurance and mitochondrial function of *Drosophila tafazzin* mutants. We further show that these beneficial effects require *sir2* and *pgc-1α*, and that overexpressing *sir2* or *pgc-1α* can rescue the exercise phenotypes of *Drosophila tafazzin* mutants. Overexpression of *pgc-1α* can also reduce the MLCL:CL ratio. Finally, we demonstrate that both muscle and neurons are key targets for future Barth syndrome therapies.

## Results

### *TAZ*^*889*^ flies have similar phenotypes to Barth patients

A new mutant allele (*TAZ*^*889*^) was generated that has the same lesion as a previously published allele (Xu et al., 2006), but with a red fluorescent protein marker knocked in to facilitate the introduction of additional transgenic elements (Fig. S1.). To confirm the new allele retained canonical Tafazzin mutant phenotypes, we examined the lipid profile and the exercise capacity of *TAZ*^*889*^ flies, since both are altered when Tafazzin function is reduced (Neuwald, 1997, Vreken et al., 2000, Xu et al., 2006, Houtkooper et al., 2009a, Schlame, 2013, Damschroder et al., 2018). The ratio of total MLCL to total CL, which is the prime diagnostic marker for BTHS (Kulik et al., 2008, Houtkooper et al., 2009b, Molenaars et al., 2021) is increased in *TAZ*^*889*^ flies (Fig. 1A., p=0.0002), with the total abundance of MLCL being higher in *TAZ*^*889*^ flies (Fig. 1B., p<0.0001). The total amount of CL is not different between *TAZ*^*889*^ flies and in control flies (*w*^*1118*^) (Fig. 1C., p=0.386). The most abundant CL species in control flies (*w*^*1118*^) were 64:4 and 66:5, and these are specifically reduced in *TAZ*^*889*^ flies (Fig. S2A.). The most abundant MLCL species to accumulate in *TAZ*^*889*^ flies was 48:3 (Fig. S2B.).

**Figure 1:**
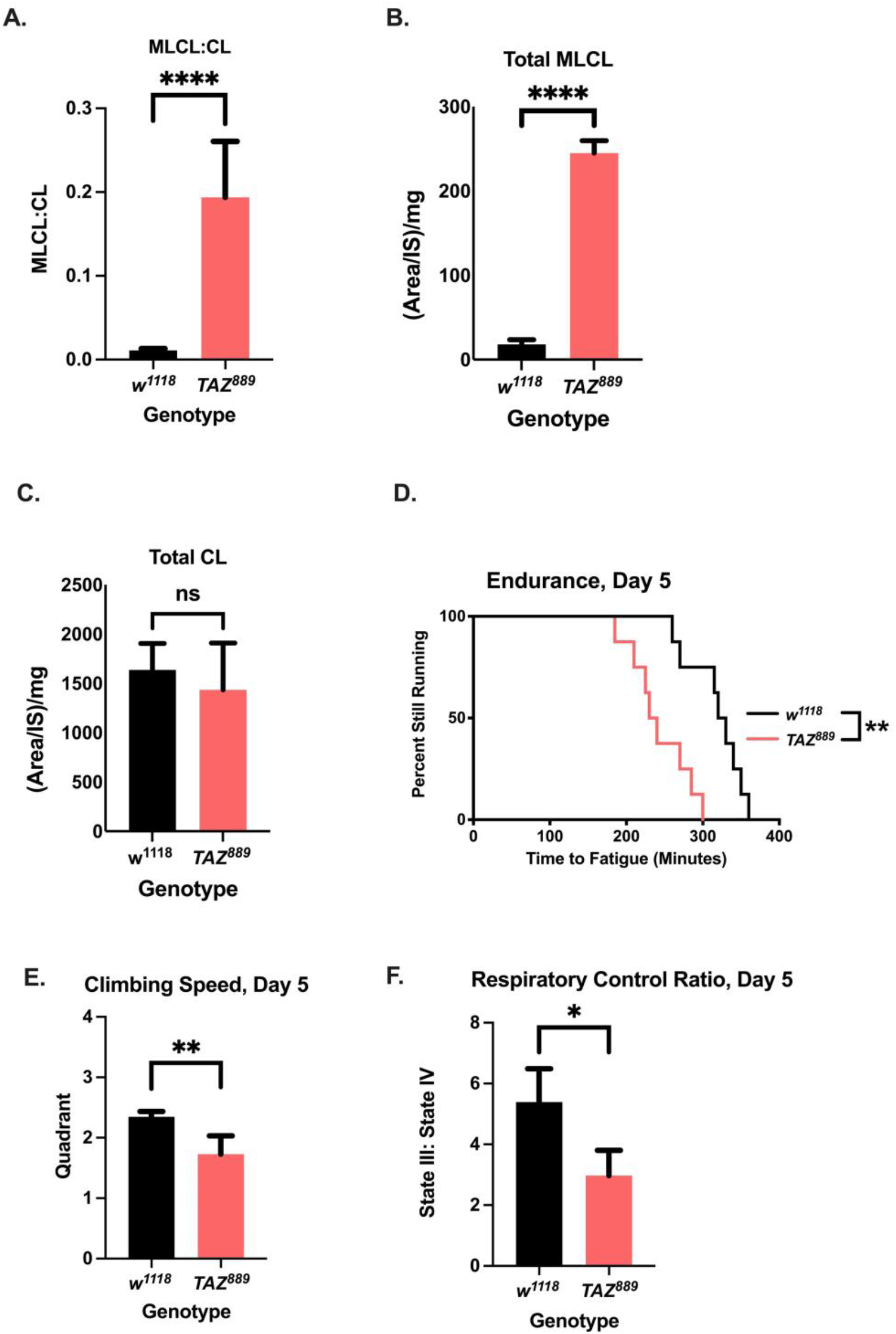
*TAZ*^*889*^ has similar phenotypes to Barth patients. (A.) The MLCL/CL ratio in *TAZ*^*889*^ flies is significantly increased (Two-tailed Student’s *t-*test, 6 biological repetitions with 6 flies per repetition). (B.,C.) *TAZ*^*889*^ flies have an increased abundance of total MLCL, but there is no difference in total CL (Two-tailed Student’s *t-*test, 6 biological repetitions with 6 flies per repetition). (D.) *TAZ*^*889*^ have reduced endurance (n=8 vials,160 flies total, log-rank analysis). (E.) *TAZ*^*889*^ flies have a reduced climbing speed (Two-tailed Student’s *t*-test, n= 100 flies, data are mean±s.d.). (F.) The respiratory control ratio of isolated mitochondria from *TAZ*^*889*^ flies is significantly reduced (Two-tailed Student’s *t*-test, six biological replicates, n=60 flies per replicate, data are mean±s.e.m.). For all experiments a p<0.05 was considered significant (*p<0.05, **p<0.01, ***p<0.001, ns=not significant).

The exercise capacity of *TAZ*^*889*^ mutants was assessed by measuring their endurance on the Power Tower (Tinkerhess et al., 2012a, Deena Damschroder, 2018). Similar to the Δ*TAZ* allele (Xu et al., 2006), *TAZ*^*889*^ flies have reduced endurance (Fig. 1D., p=0.0014) and reduced climbing speed (Fig. 1E., p=0.0078) (Damschroder et al., 2018). To test mitochondrial function in mutants, the respiratory control ratio (RCR) of isolated mitochondria was measured. The RCR is the ratio of state III respiration (ADP stimulated) to state IV respiration (oligomycin stimulated) and reflects general mitochondrial health (Gonzalvez et al., 2013). Mutants have a reduced RCR, indicating their mitochondria have reduced efficiency of mitochondrial coupling (Fig. 1F., p=0.038). Taken together, these results demonstrate that the *TAZ*^*889*^ allele retains stereotypical phenotypes of reduced TAZ function.

### *Nicotinamide riboside supplementation rescues the exercise capacity of TAZ*^*889*^ flies

*TAZ*^*889*^ flies have a higher abundance of NAD^+^ and NADH than control flies from the same genetic background (*w*^*1118*^) (Fig. 2A., Tukey *post-hoc* test, p=0.0017), while the NAD^+^:NADH ratio is significantly reduced in the mutants (Fig. 2C., Tukey *post-hoc* test, p=0.0002). NR supplementation at various concentrations caused improvements to endurance, but 1 mM generated the largest improvement to endurance (Fig. S3A., p<0.0001). The NR concentration in the food did not affect the feeding rate (Fig. S3B., p=0.08). NR supplementation did not significantly change the levels of NAD^+^, NADH, or the NAD^+^:NADH ratio between NR-fed mutants and vehicle fed mutants (Fig. 2A., Tukey *post-hoc* test, p=0.671, 2B, Tukey *post-hoc* test p=0.722, 2C Tukey *post-hoc* test, p=0.991).

**Figure 2:**
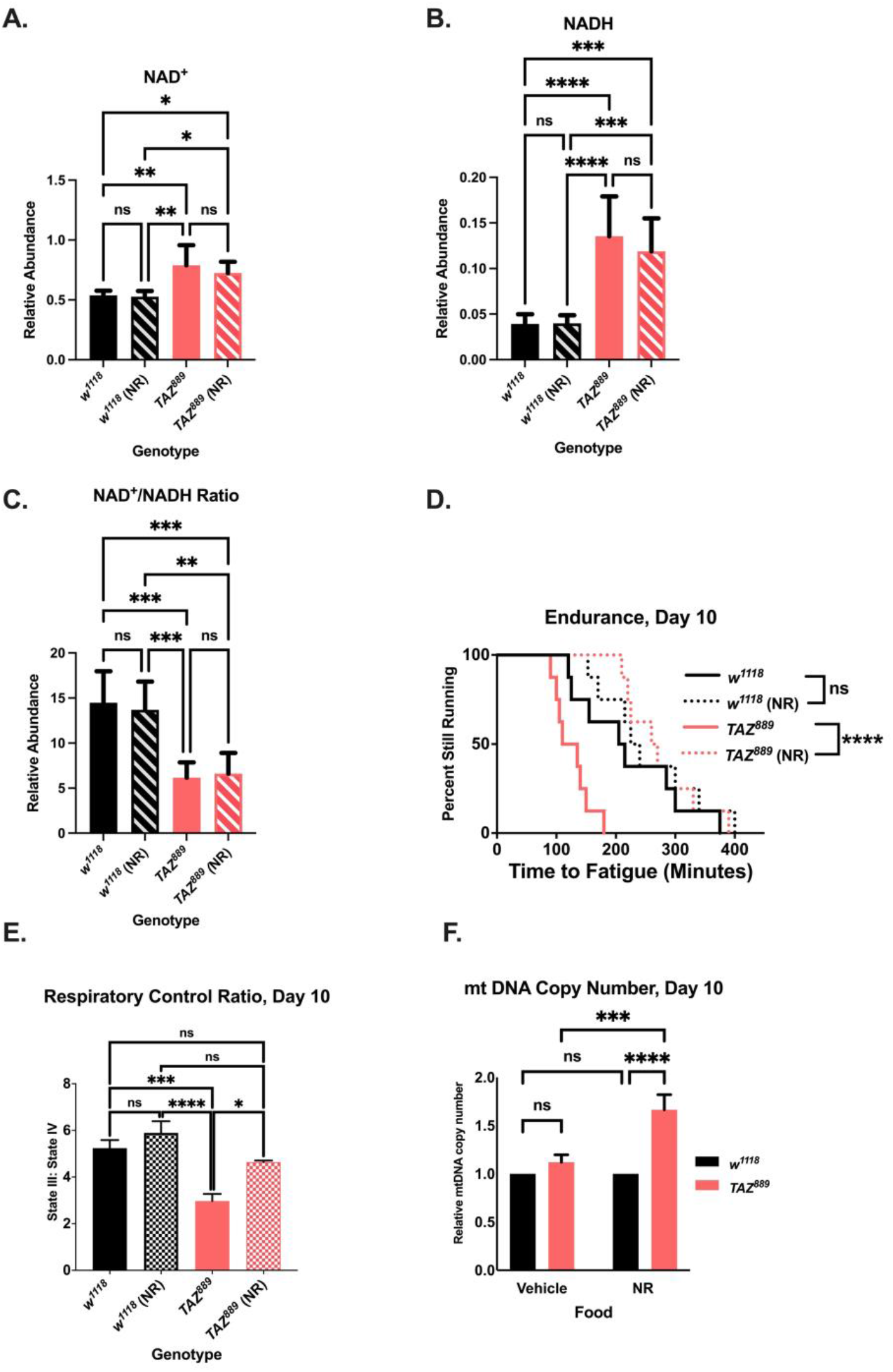
NR supplementation provides benefits to *TAZ* mutants. (A.,B.) The abundance of NAD^+^ and NADH was measured by mass spectrometry and normalized to total protein levels (6 biological repetitions with 6 flies per repetition, data are mean±s.d., 2-way ANOVA, genotype effect, p<0.0001, Tukey *post-hoc* test). (C.) The NAD^+^:NADH ratio was calculated from the abundance for those molecules (data are mean±s.d., 2-way ANOVA, genotype effect, p<0.0001, Tukey *post-hoc* test). (D.,E.) NR supplementation restores the endurance of *TAZ*^*889*^ flies (log-rank analysis, n=8 vials, 20 flies per vial) and the RCR (6 biological replicates, data are mean±s.e.m, n=60 per replicate, genotype effect, p<0.0001, NR effect, p=0.0029,Tukey *post-hoc* test). (F.) At age day 10, the relative mt DNA copy number is not different between *TAZ*^*889*^ and control flies, but NR supplementation increases mt DNA in *TAZ*^*889*^ flies (n= 3 biological replicates, data are mean±s.d., 2-way ANOVA, genotype effect, p<0.0001, NR effect, p=0.0006, Tukey *post-hoc* test). A p-value of less than 0.05 was considered significant. *p<.05, **p<.01, ***p<.001, ****p<0.0001, ns=not significant

However, the endurance of NR-fed mutants increases after 5 days of supplementation (Fig. 2D., p<0.0001), while supplementation for 3 days does not improve endurance (Fig. S3C., p=0.487). The RCR of NR-fed mutants is also rescued to control levels (Fig. 2E., Tukey *post-hoc* test, p=0.611). Supplementation provided no benefit to the control line in endurance or RCR (Fig. 2E., Tukey *post-hoc* test, p=0.544). To determine if there was a difference in mitochondrial number, the mtDNA copy number was measured using the relative gene expression of a mitochondrial gene, lrRNA, to nuclear gene, rp2 (Correa et al., 2012). NR-fed mutants have an increase in mitochondrial DNA copy number relative to vehicle-fed mutants (Fig. 2F., Tukey *post-hoc* test, p=0.0003), while NR-fed control flies do not have more mitochondria relative to vehicle-fed controls (Fig. 2F., Tukey *post-hoc* test. p=0.978). Thus, NR supplementation to mutants can restore endurance, improve mitochondrial function, and increase mitochondrial number, without measurably changing total NAD^+^ and NADH content or the NAD^+^:NADH ratio.

A possible reason that the levels of NAD^+^, NADH and the NAD^+:^NADH ratio did not change with supplementation is that excess NAD^+^ is rapidly utilized to induce the rescue phenotype (Fang et al., 2017). A possible consumer of NAD^+^ that could be responsible for the rescue phenotype is *sir2*, which is a NAD^+^ -dependent deacetylase (Stein and Imai, 2012). Sir2 has many targets, including the well-studied exercise protein PGC-1α (Baar et al., 2002, Rodgers et al., 2005, Handschin et al., 2007, Olesen et al., 2010, Tinkerhess et al., 2012b). Therefore, we hypothesized that *sir2* and *pgc-1α* are required for the rescue phenotypes observed with NR supplementation. Using mifepristone inducible gene-switch lines, we induced whole-body changes to gene expression in an adult specific manner. Gene expression was confirmed using qRT-PCR (Fig. S4). When *sir2* is knocked down in *TAZ*^*889*^ flies, NR no longer increases endurance or improves RCR (Fig. 3A., p=0.432, Fig. 3B., 2-way ANOVA, Tukey *post-hoc* test, p=0.988). Likewise, when *spargel* is mutated in *TAZ*^*889*^ flies, NR supplementation cannot increase the endurance (Fig. 3C., p=0.25) or the RCR (Fig. 3D., Tukey *post-hoc* test, p=0.936). Taken together, these results confirm that *sir2* and *pgc-1α* are required for NR supplementation to increase endurance or RCR.

**Figure 3:**
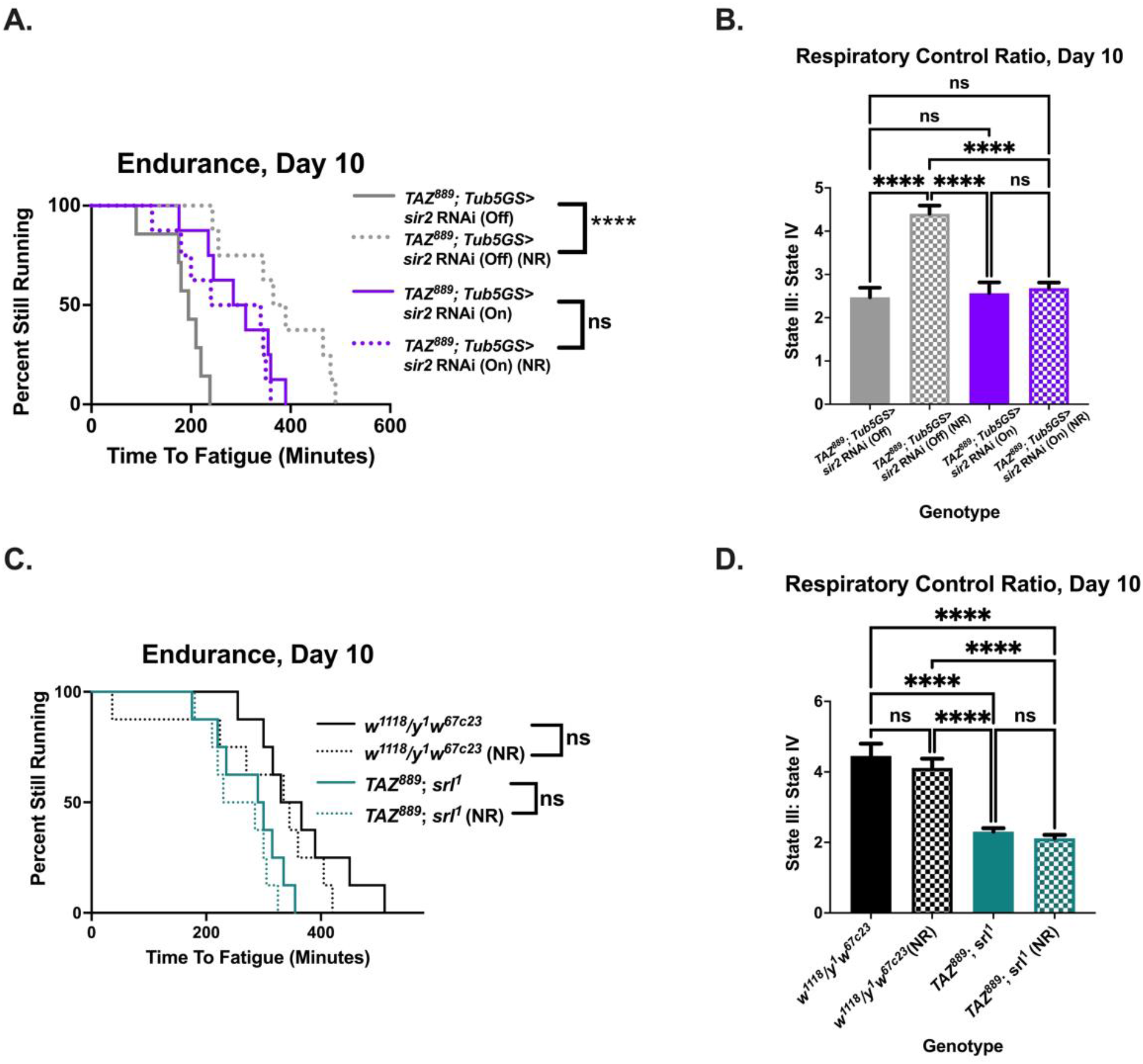
The beneficial effects of NR are *sir2* and *srl* dependent. (A., B.) When *sir2* is knocked down in *TAZ*^*889*^ flies, NR supplementation does not improve their endurance (*TAZ*^*889*^;*Tub5GS>uas-sir2* RNAi On) (log-rank analysis, n=8 vials, 20 flies per vial). The RCR is not restored by NR supplementation when *sir2* is knocked down in *TAZ*^*889*^ flies (6 biological replicates, data are mean±s.e.m, n=60 flies per biological replicate, 2-way ANOVA, genotype effect, p=0.002, Tukey *post-hoc* test). (C.) When the *Drosophila pgc-1α* homolog *spargel* is mutated (*srl*^*1*^) in *TAZ*^*889*^ flies, NR feeding does not improve the endurance of *TAZ*^*889*^ flies (log-rank analysis, n=8 vials, 20 flies per vial). (D.) The RCR of the double mutant (*TAZ*^*889*^;*srl*^*1*^) is not improved after NR supplementation (6 biological replicates, data are mean±s.e.m, n=60 flies per replicate, 2-way ANOVA, genotype effect, p<0.0001, Tukey *post-hoc* test). A p-value <0.05 was considered significant (*p<0.05, **p<.01, ***p<.001, ****p<0.0001, ns=not significant). On=mifepristone induced gene

### Overexpressing *sir2* or *pgc-1α* is sufficient to restore the endurance of *TAZ*^*889*^ flies

We next hypothesized that overexpressing *sir2* or *pgc-1α* would be sufficient to restore endurance to *TAZ*^*889*^ flies. The endurance of mutants is increased when *sir2* is overexpressed (Fig. 4A., p<0.0001) and there is no additive effect with NR supplementation (Fig. 4A., p=0.093). The RCR of mutants is also improved with *sir2* overexpression (Fig. 4B.,Tukey *post-hoc* test, p<0.0001), but NR supplementation does not provide further improvements to *sir2* rescue flies (Fig. 4B., Tukey *post-hoc* test, p=0.793). Overexpression of *spargel* in mutants produces similar results, causing an increase to endurance (Fig. 4C., p<0.0001) and the RCR (Fig. 4D., Tukey *post-hoc* test, p=0.0035). There were no additive improvements to the endurance (Fig. 4C. p=0.23) or the RCR with NR supplementation (Fig. 4D., Tukey *post-hoc* test, p=0.773).

**Figure 4:**
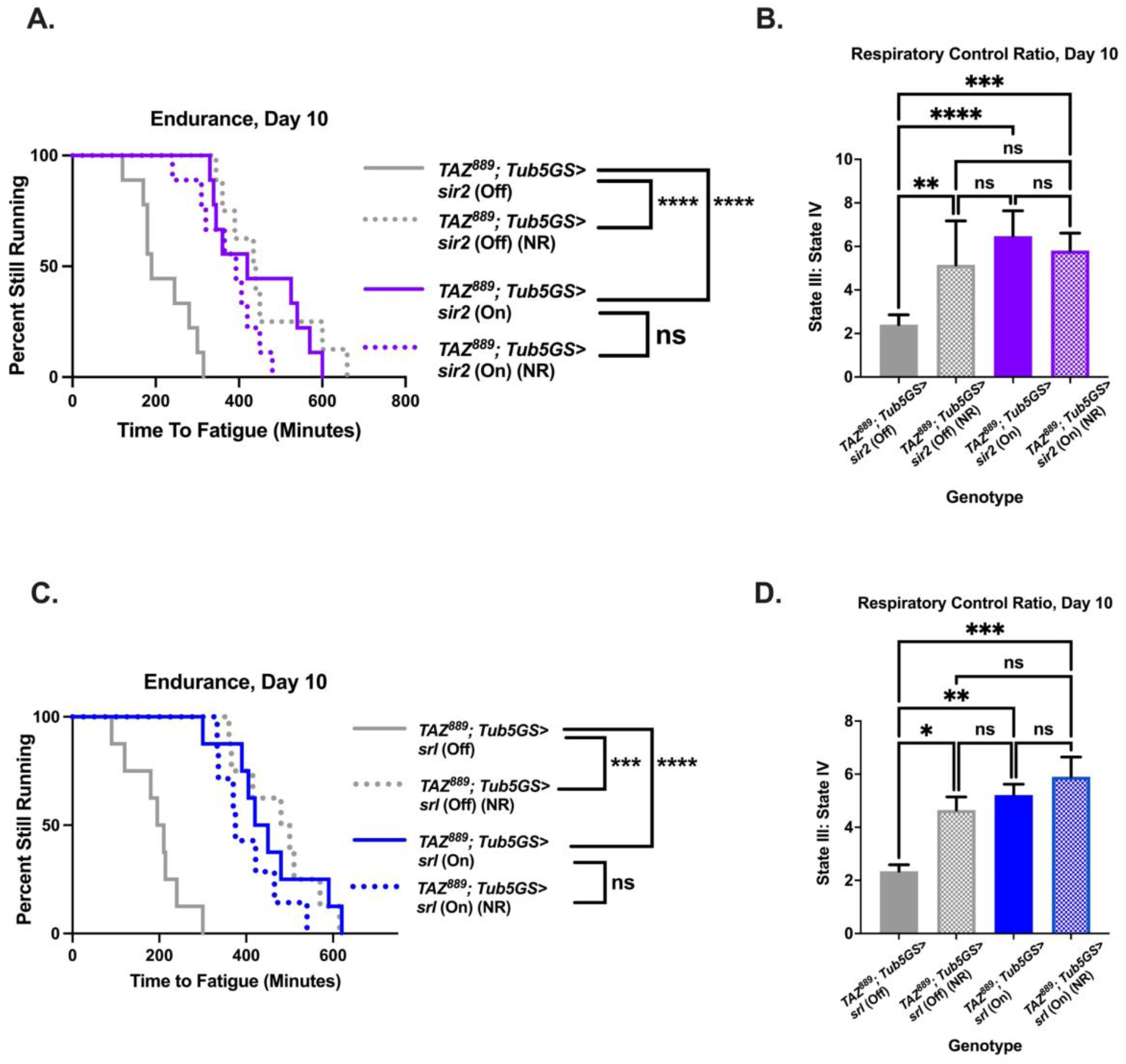
Overexpression of *sir2* or *srl* is sufficient to improve the exercise capacity of *TAZ*^*889*^ flies. (A.,B.) Overexpressing *sir2* in *TAZ*^*889*^ flies (*TAZ*^*889*^;*Tub5GS> uas-sir2* On) increases their endurance (n= 8 vials, 160 flies total, pair-wise log rank analysis) and improves the RCR (6 biological replicates, data are mean±s.e.m, n=60 flies per replicate, 2-way ANOVA, genotype effect, p=0.0002, Tukey *post-hoc* test). NR feeding does not provide further improvements to these phenotypes. (C.,D.) The endurance (n= 8 vials, 160 flies total, pair-wise log rank analysis) and the RCR (6 biological replicates, data are mean±s.e.m, n=60 flies per replicate, 2-way ANOVA, genotype effect, p=0.0002, NR effect, p=0.045, Tukey *post-hoc* test) of *TAZ*^*889*^ flies overexpressing *spargel* (*TAZ*^*889*^;*Tub5GS>uas-srl* On) are increased, but supplementation with NR does not provide any additive benefits. p<0.05, **p<.01, ***p<.001, ****p<0.0001, ns=not significant, On=mifepristone induced gene

Additionally, no additive effects were observed with NR supplementation to control flies overexpressing *sir2* or *spargel* (Fig. S5).

### Overexpressing *pgc-1α* in *TAZ*^*889*^ flies improves the NAD^+^:NADH ratio, increases mitochondrial DNA, and reduces the MLCL:CL ratio

To determine if overexpression of *spargel* in *TAZ*^*889*^ flies influenced the NAD^+^:NADH ratio, we measured the relative abundance of NAD^+^ and NADH. There was no difference in the abundance of NAD^+^ (Fig. 5A.). *TAZ*^*889*^ flies overexpressing *spargel* have a lower NADH abundance than mutants (Fig. 5B., p<0.0001) or *w*^*1118*^ flies (Fig. 5B., p<0.0001), resulting in a higher NAD^+^:NADH ratio (Fig. 5C., p<0.0001). The reduction in NADH is not due to increased lactate production (Fig. 5D.), suggesting a possible effect at the mitochondrial level. To determine if there was an increase in mitochondrial biogenesis, the mtDNA copy number was measured. *TAZ*^*889*^ flies overexpressing *spargel* have an increase in mtDNA copy number (Fig. 5E, p=0.0012), but there are no differences in the glutathione (GSH) to glutathione disulfide (GSSG) ratio (Fig. 5F., p=0.316), indicating that there is not an increase of oxidative stress. The MLCL:CL ratio in *TAZ*^*889*^ flies overexpressing *spargel* is significantly reduced (Fig. 5G., p=0.032). There is no change in the total amount of MLCL between *TAZ*^*889*^ flies overexpressing *spargel* and mutants (Fig. 5H., p=0.945), but *TAZ*^*889*^ flies overexpressing *spargel* have an increase in total CL relative to mutants (Fig. 5I., p=0.023).

**Figure 5:**
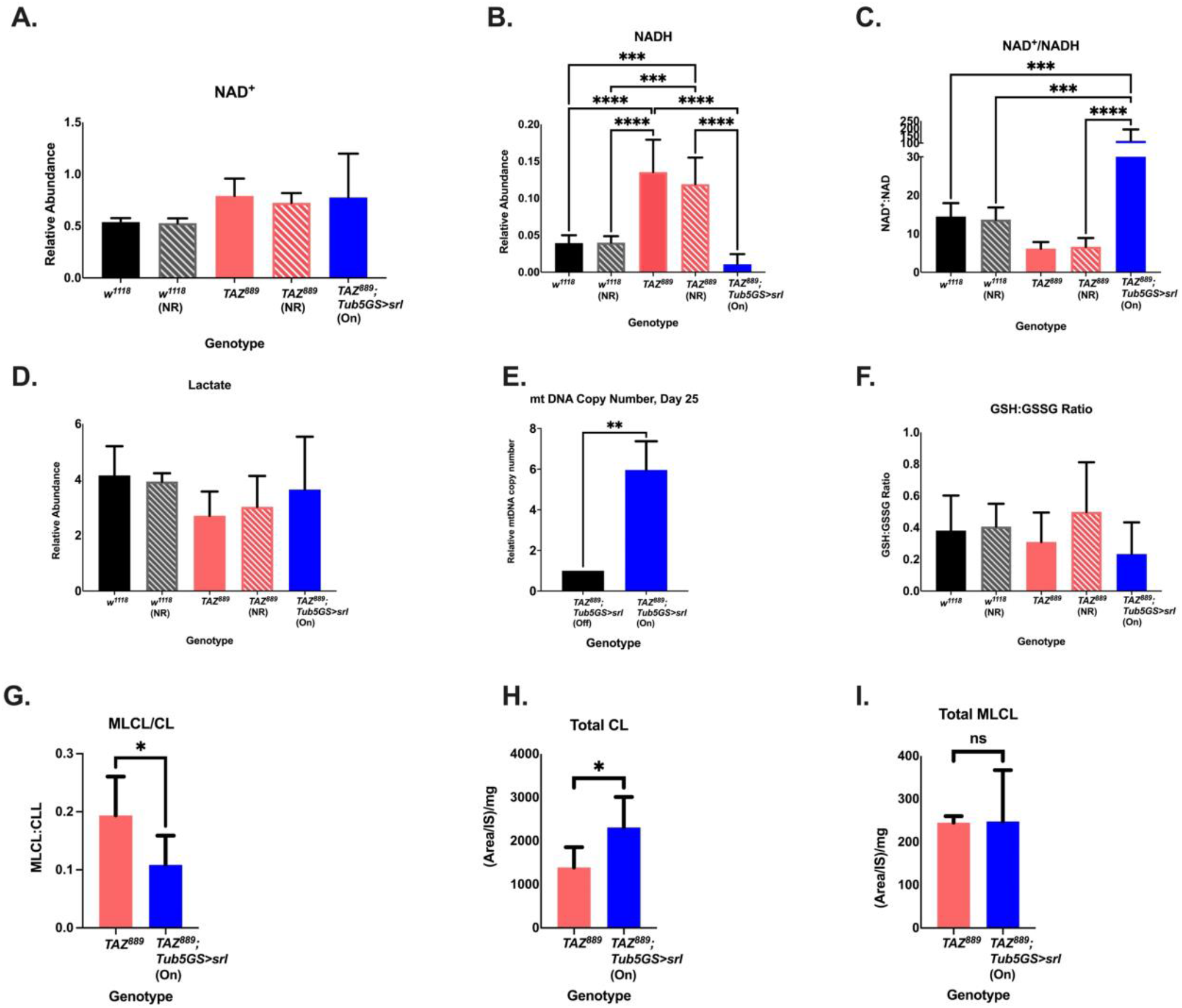
Overexpressing *spargel* in *TAZ*^*889*^ flies improves the NAD^+^:NADH ratio, increases the mitochondrial DNA copy number and reduces the MLCL:CL ratio. All measurements were performed on flies age day 10, exception the measurement of mtDNA copy. The abundance of NAD^+^ and NADH was measured by mass spectrometry and normalized to total protein levels (6 biological repetitions with 10 flies per repetition, data are mean±s.d., one-way ANOVA, Tukey Post-Hoc test). (A.) There is no significant difference in the relative abundance of NAD^+^ (One-way ANOVA, p=0.096). (B.) Overexpressing *spargel* in *TAZ*^*889*^ flies significantly reduces NADH relative to all other groups, (C.) consequently increasing the NAD^+^:NADH ratio. (D.) There is no difference in lactate production between experimental groups (one-way ANOVA, p=0.194). (E.) mtDNA copy number represents the relative expression of the mitochondrial gene lrRNA to the nuclear gene rp2. At age day 25, *TAZ*^*889*^ flies overexpressing *spargel* have a higher mtDNA copy relative to *TAZ*^*889*^ flies (n= 3 biological replicates, data are mean±s.d., Student’s *t-* test). (F.) The ratio of glutathione (GSH) to glutathione disulfide (GSSG) is not significantly different between groups (one-way ANOVA, p=0.316 (G.) The MLCL:CL ratio of TAZ889 flies overexpressing spargel is significantly reduced relative to TAZ889 flies (Two-tailed Student’s t-test, three biological replicates, n=6 flies per biological replicate, data are mean±s.d.). (H.,I.) The reduced MLCL:CL ratio is not due to a reduced total MLCL amount (Two-tailed Student’s t-test, three biological replicates, n=6 flies per biological replicate, data are mean±s.d., p=0.954), but to an increased total CL amount (Two-tailed Student’s t-test, three biological replicates, n=6 flies per biological replicate, data are mean±s.d.). *p<.05, **p<.01, ***p<.001, ****p<0.0001, ns= not significant, On=mifepristone induced gene

### Tafazzin is required in both muscle and neurons for normal endurance

Patients with Barth syndrome often display exercise intolerance due to their skeletal myopathy, low muscle tone, and cardiomyopathy (Barth et al., 1983, Spencer et al., 2006, Thompson et al., 2016). However, the tissue specific requirements for *tafazzin* for normal exercise capacity have not been rigorously investigated. We wanted to identify the key tissues responsible for the exercise intolerance of *tafazzin* mutants as a first step to investigating if tissue specific expression of *sir2* or *spargel* could rescue endurance and mitochondrial function. Prior to tissue specific knockdown experiments, a whole-body driver was used to validate the RNAi line and, like the genomic mutant, ubiquitous knock down of *tafazzin* caused reduced endurance (Fig. S6A., p=0.0006), climbing speed (Fig. S6B., p=0.012) and RCR (Fig. S6C., p<0.0001).

In the fly, *tafazzin* is required in both muscle (Fig. 6A., p=0.004) and neurons (Fig. 6B., p=0.016) for normal endurance, while knocking down *tafazzin* in the heart or fat body causes no significant effect on endurance (Fig.S6D., p=0.113, Fig. S6E., p=0.2134). Climbing speed is also significantly reduced in both the muscle-specific and neuron-specific knockdown (Fig. 6C., p=0.0149, Fig. 6D., p=0.013). To examine the mitochondrial function of the tissue-specific knockdown flies, we isolated mitochondria from heads, which are enriched for neurons, and from the thoraces, which are enriched with muscle tissue. The RCR of mitochondria from the thoraces is reduced in the muscle-specific *tafazzin* knockdown (Fig. 6E., Tukey *post-hoc* test, p=0.002) and the RCR of mitochondria from heads was reduced in the neuron-specific *tafazzin* knockdown (Fig. 6F., Tukey *post-hoc* test p=0.0001). To test the specificity of the tissue-specific drivers, the gene expression of *tafazzin* was measured in the heads and thoraces of these tissue-specific knockdown flies. Muscle-specific *tafazzin* knockdown flies had reduced *tafazzin* expression in their thoraces relative to their heads and the neuronal-specific *tafazzin* knockdown flies had reduced *tafazzin* expression in their heads relative to their thoraces (Fig. S4B.), validating the specificity of the tissue drivers.

**Figure 6:**
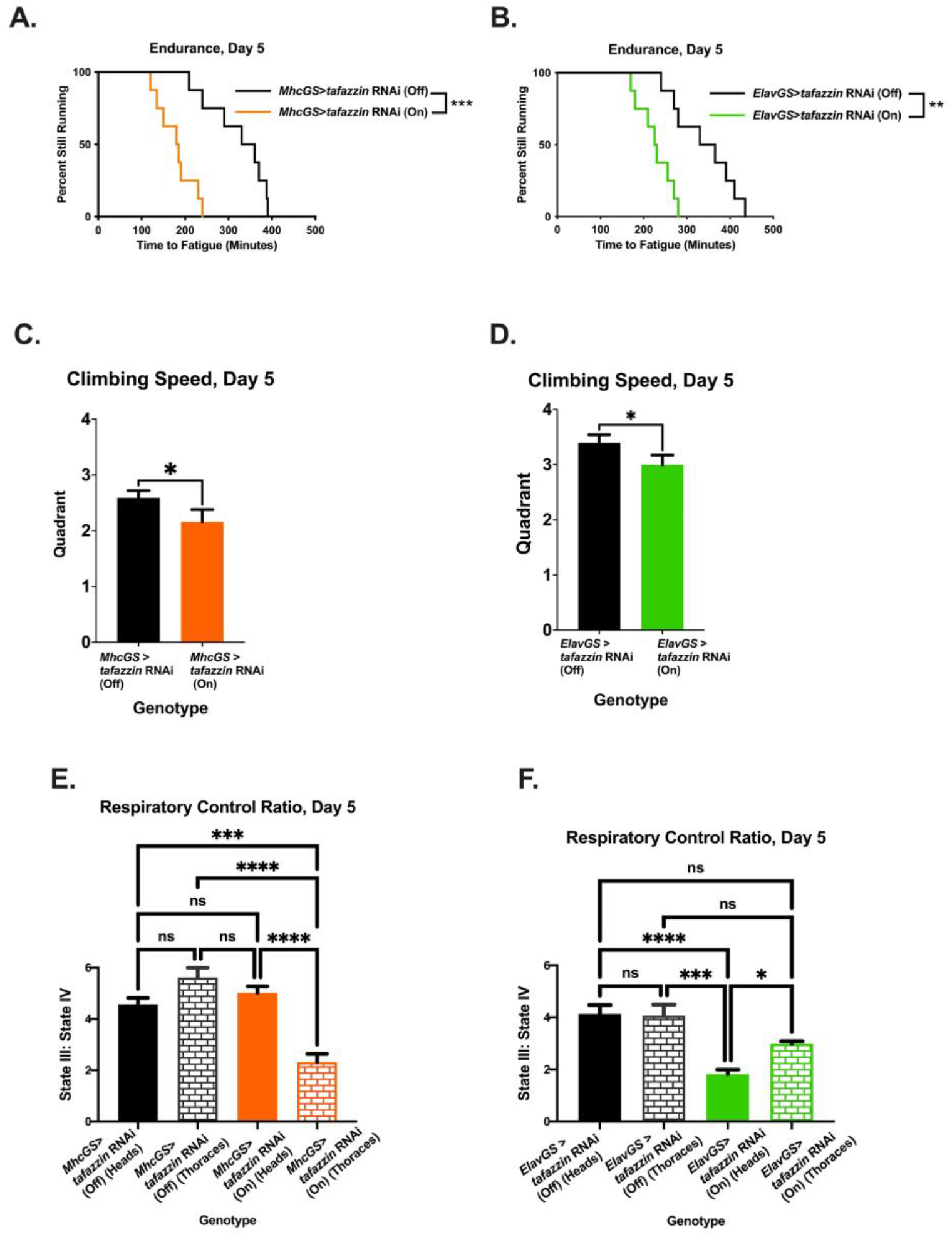
*tafazzin* is required in either muscle or neurons for normal exercise capacity. (A.,B.) Knocking down *tafazzin* in muscle or neuronal tissue reduces endurance (log-rank analysis, n=8 vials, 20 flies per vial). (C.,D.) These tissue specific knockdown flies have a reduced climbing speed (Two-tailed Student’s *t*-test, n= 100 flies, data are mean±s.d). (E., F.) The RCR of the muscle enriched thoraces (2-way ANOVA, genotype effect, p=0.0002, tissue effect, p=0.013,Tukey *post-hoc* test) or the neuronal enriched heads (2-way ANOVA, genotype effect, p<0.0001, tissue effect, p<0.0001, Tukey *post-hoc* test) are reduced relative to background controls and counter body part (data averages of 6 biological replicates ±s.e.m, n=60 flies per replicate). p<0.05, **p<.01, ***p<.001, ****p<0.0001, ns=not significant, On=mifepristone induced gene expression

We next tested if overexpressing *tafazzin* in the muscle or neuronal tissue of *TAZ*^*889*^ flies would be beneficial. The endurance of the muscle-specific and the neuronal-specific rescue of *tafazzin* is not significantly improved at age day 5 (Fig. S7A., p=0.679, Fig. S7B., p=0.378), but by age day 12 the endurance is higher (Fig. 7A., p<0.0001 & Fig. 7B., p<0.0001). The climbing speed of the muscle-specific rescue and the neuronal-specific rescue is also improved at age day 12 (Fig. 7C., p=0.0013, Fig. 7D., p=0.0023) along with the RCR from the rescue tissues (Fig. 7E., Tukey *post-hoc* test, p<0.0001, Fig. 7F, Tukey *post-hoc* test, p<0.0001). The relative abundance of *tafazzin* is not significantly different between day 5 and day 12 for the muscle and the neuronal specific rescue (Fig. S4C., p=0.982, p=0.998), so the rescue phenotype at age day 12 is not due to increased *tafazzin* expression. These results confirm muscle and neurons as a key target for future Barth treatments.

**Figure 7:**
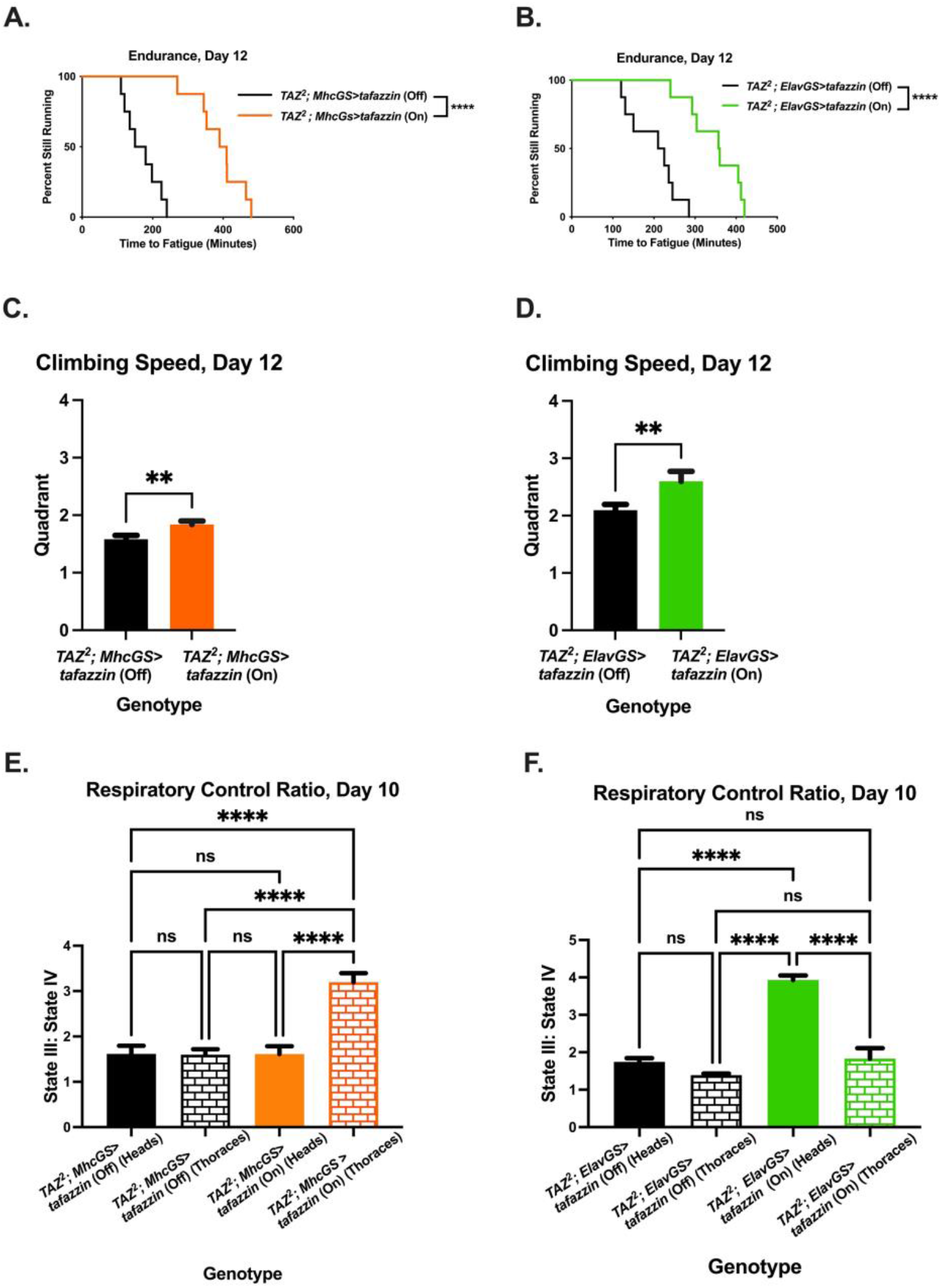
Restoring *tafazzin* in either muscle or neurons is sufficient to increase exercise capacity. (A.,B.) When *tafazzin* is restored in *TAZ*^*2*^ flies in either muscles or neurons, the endurance is significantly increased by day 12 (log-rank analysis, n=8 vials, 20 flies per vial) (C.,D.) The climbing speed is also improved by day 12 (Two-tailed Student’s *t*-test, n= 100 flies, data are mean±s.d). (E.,F.) The RCR is increased in mitochondria isolated from the thoraxes of the muscle specific rescue (2-way ANOVA, genotype effect, p=0.0001, tissue effect, p=0.0002, Tukey *post-hoc* test) and in mitochondria isolated from the heads of the neuronal specific rescue (data averages of 6 biological replicates ±s.e.m, n=60 per replicate, 2-way ANOVA, genotype effect, p<0.0001, tissue effect, p<0.0001, Tukey *post-hoc* test). A p-value of less than 0.05 was considered significant. *p<.05, **p<.01, ***p<.001, ****p<0.0001, ns= not significant, On=mifepristone induced gene expression

### Tissue specific overexpression of *sir2* and *pgc-1α* improves endurance of *TAZ*^*889*^ flies

Overexpressing *sir2* or *spargel* ubiquitously in *TAZ*^*889*^ flies provides substantial benefits (Fig. 3). We hypothesized that overexpressing these genes in just muscle or neurons of mutants could be sufficient to rescue their endurance. *sir2* overexpression in the muscle or neurons of mutants rescues their endurance (Fig. 8A., p<0.0001, Fig. 8B., p=0.0034). When *sir2* is expressed in muscle, mitochondrial RCR is increased in thoraces but not in the heads (Fig. 8C., Tukey *post-hoc* test, p<0.0001). When *sir2* is expressed in neurons, mitochondrial RCR is increased in heads but not thoraces (Fig. 8D., Tukey *post-hoc* test, p<0.0001), which is consistent with tissue-autonomous effect of Sir2.

**Figure 8:**
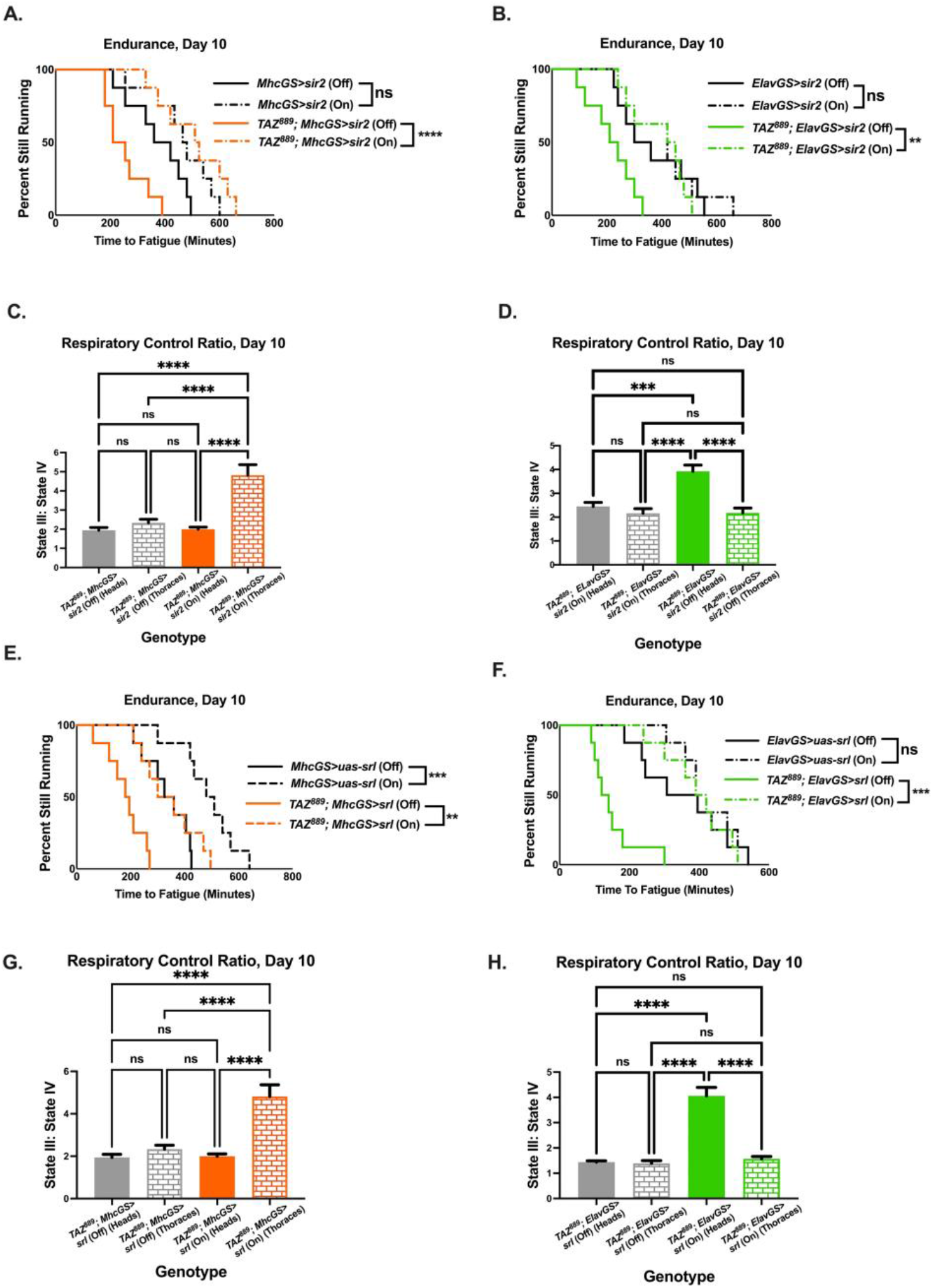
Tissue specific expression of *sir2* or *spargel* causes beneficial phenotypes in *TAZ*^*889*^ flies. (A., B.) Overexpressing of *sir2* in muscle or neurons of *TAZ*^*889*^ flies restores their endurance (log-rank analysis, n=8 vials, 20 flies per vial). (C.) Muscle-specific overexpression of *sir2* restores the RCR of mitochondria isolated from thoraces (2-way ANOVA, genotype effect, p<0.0001, tissue effect, p=0.0005, Tukey *post-hoc* test) (data averages of 6 biological replicates ±s.e.m, n=60 flies per replicae). (D.) Neuronal-specific overexpression of *sir2* improves the RCR of mitochondria isolated from heads (2-way ANOVA, genotype effect, p=0.0001, tissue effect, p=0.0021, Tukey *post-hoc* test) (data averages of 6 biological replicates ±s.e.m, n=60 flies per replicate). (E., F.) *srl* overexpressed in muscle or neurons improves the endurance of *TAZ*^*889*^ flies (log-rank analysis, n=8 vials, 20 flies per vial). (G.) Muscle-specific overexpression of *srl* increase the RCR of mitochondria from thoraces (2-way ANOVA, genotype effect, p<0.0001, tissue effect, p=0.0005, Tukey *post-hoc* test) (data averages of 6 biological replicates ±s.e.m, n=60 flies per replicate). (H.) Overexpressing *srl* in neurons improves the RCR of mitohicdnira isolated from heads (2-way ANOVA, genotype effect, p<0.0001, tissue effect, p<0.0001, Tukey *post-hoc* test) (data averages of 6 biological replicates ±s.e.m, n=60 flies per replicate. A p-value of less than 0.05 was considered significant. *p<.05, **p<.01, ***p<.001, ****p<0.0001, ns=not significant, On=mifepristone induced gene expression

Overexpressing *spargel* has a similar effect in both tissues. The endurance of *TAZ*^*889*^ flies is improved when *spargel* is expressed in muscle (Fig. 8E., p=0.0017) or neurons (Fig. 8F., p=0.0002). The RCR of mitochondria isolated from the thoraces of *TAZ*^*889*^ flies overexpressing *spargel* in their muscle is improved while mitochondria isolated from the heads is not (Fig. 8G., Tukey *post-hoc* test, p<0.0001). By contrast, overexpressing *spargel* pan-neuronally in *TAZ*^*889*^ flies improves the RCR of mitochondria isolated from heads, but not thoraces (Fig. 8H., Tukey *post-hoc* test, p<0.0001), which is again consistent with a tissue-autonomous effect.

## Discussion

When cardiolipin’s remodeling process is disturbed by *tafazzin* mutations, the structural changes to the mitochondrial membrane cause reduced mitochondrial function and Barth syndrome (Neustein et al., 1979, Schlame et al., 2002). Although there is currently no cure for Barth syndrome, *tafazzin* replacement therapy using adeno-associated virus (AAV) vector shows promising results in *tafazzin* knock-down mice, including improved mitochondrial structure, mitochondrial respiration, and heart function (Suzuki-Hatano et al., 2019). However, AVV gene therapy as a cure for Barth syndrome does not yet have FDA approval and not all patients may have access to facilities capable of administering the treatment. Therefore, it is important for other therapeutic avenues to be examined.

Declining NAD^+^ abundance is a hallmark of mitochondrial diseases (Srivastava, 2016). Restoring NAD^+^ content in various disease models provides numerous benefits, including improved mitochondrial function (Yoshino et al., 2011, Canto et al., 2012, Schondorf et al., 2018). In *TAZ*^*889*^ flies the NAD^+^:NADH ratio is reduced. We demonstrate for the first time that NR supplementation is sufficient to rescue the exercise capacity and mitochondrial coupling of *TAZ*^*889*^ flies, despite causing no evident accumulation of NAD^+^ or changes in the NAD^+^:NADH ratio. NR supplementation provided no benefits to control flies, which supports the idea that NR supplementation is compensating for a deficit specific to *TAZ*^*889*^ flies and is not a general booster to endurance.

Like our results, one study using the genetic mitochondrial disease mouse model (*Sco2* knockout/knockin model) found that NR supplementation improved the mutant’s exercise capacity while their control group gained no benefits (Cerutti et al., 2014). Multiple studies demonstrate that benefits from NR or NA supplementation work though Sir2 (Cerutti et al., 2014, Khan et al., 2014, Pirinen et al., 2020), further supporting the findings in this study. Others report that NR supplementation increases the abundance of NAD^+^ in various model systems including cells (Schondorf et al., 2018) and mice (Canto et al., 2012, Trammell et al., 2016, Schondorf et al., 2018, Yang et al., 2020). The tissue type the NAD^+^ measurements originated from is an important consideration when examining those studies. In mice, NAD^+^ flux experiments demonstrate that tissues metabolize NAD^+^ at different rates (Liu et al., 2018) and the natural abundance of NAD^+^ is tissue specific (Canto et al., 2012, Liu et al., 2018). Furthermore, NR’s ability to increase NAD^+^ concentration appears to be tissue specific, with an increase in NAD^+^ after NR supplementation in mice identified in liver, skeletal muscle, or brown adipose tissue, but not in white adipose tissue or brain (Canto et al., 2012). In humans, NR supplementation is reported to increase NAD^+^ concentrations in the peripheral blood mononuclear cells of healthy older participants (Trammell et al., 2016, Martens et al., 2018), but not in the skeletal muscle of young (Stocks et al., 2021), or obese (Remie et al., 2020) subjects. Together, these studies indicate that tissue type influences the baseline NAD^+^ abundance and possibly the ability for NR to increase NAD^+^ concentrations.

The metabolomics analysis performed in this study was on whole flies, so it is not possible to determine the tissue or cellular origins of metabolites. By pooling all the tissues together, it is possible that tissue-specific changes in the abundance of NAD^+^ and its precursors due to NR supplementation were not detected.

NAD^+^ exists in three intracellular compartments: cytosolic, nuclear, and mitochondrial (Cambronne and Kraus, 2020, Amjad et al., 2021). Elevated NAD^+^ abundance within these pools can stimulate key NAD^+^ consuming enzymes located within that compartment (Cambronne and Kraus, 2020, Amjad et al., 2021). NR supplementation stimulates SIRT1 and SIRT3 in mice (Canto et al., 2012, Brown et al., 2014), which are located in the nucleus and mitochondria respectively (Onyango et al., 2002, Michishita et al., 2005, Zakhary et al., 2010). Therefore, NR supplementation likely increases the abundance of NAD^+^ in the nuclear and mitochondrial compartment (Canto et al., 2012, Cambronne and Kraus, 2020, Mehmel et al., 2020).

We demonstrate that Sir2 (homologous to mammalian SIRT1) is required for NR’s beneficial effects in *tafazzin* mutants. NR supplementation is likely increasing the abundance of NAD^+^ in the nuclear compartment. Measuring NAD^+^ abundance *in vivo* and without disturbing the physiological environment within the compartments, is complicated and current techniques are unable to distinguish between bound and free NAD^+^ (Cambronne, 2020). Therefore, in this study, we did not measure the abundance of NAD^+^ within these intracellular compartments. Future work may help illuminate the dynamic interactions between the different NAD^+^ pools and SIRT1 activity in the context of BTHS.

Metabolomics data shows that overexpressing *spargel* in *TAZ*^*889*^ flies affects the NAD^+^:NADH ratio in whole flies more than NR supplementation. The improved NAD^+^:NADH ratio is likely due to an increase in NADH oxidation since the abundance of NADH returns to wild-type levels while there was no difference in the abundance of NAD^+^. This suggests that NADH is being more efficiently utilized in *TAZ*^*889*^ flies overexpressing *spargel* relative to *TAZ*^*889*^ flies fed NR.

NADH is oxidized by several enzymes including lactate dehydrogenase to replenish NAD^+^ during glycolysis and by complex I of the mitochondrial respiratory chain (Berrisford and Sazanov, 2009, Sazanov, 2015). There is no increase in NADH oxidation by lactate dehydrogenase, since there is no increase in lactate production between the experimental groups. Overexpression of *pgc-1α* increases the number of mitochondria (Lehman et al., 2000, Tiefenbock et al., 2010) and the proteins of the electron transport chain (Wu et al., 1999). Therefore, due to the increase in mitochondrial number, and probable increase in complex I proteins, it is likely that more NADH is being oxidized by mitochondria in *TAZ*^*889*^ flies overexpressing *spargel*, resulting in lower NADH content and improved mitochondrial coupling.

Disrupted cardiolipin remodeling is a key phenotype of Barth syndrome (Vreken et al., 2000). The most abundant CL species in control flies (64:4 and 66:5) are reduced in *TAZ*^*889*^ flies. The CL species 64:4 and 66:5 is most likely comprised of the smaller CL species 16:1, which was also found to be the most abundant CL in wild-type flies (Xu, 2006). The MLCL:CL ratio is also important in the context of Barth syndrome since patients with a lower MLCL:CL ratio tend to have less severe phenotypes (Bowron et al., 2015). Coinciding with human data (Bowron et al., 2015), *TAZ*^*889*^ flies overexpressing *spargel* have a reduced MLCL:CL ratio and their exercise capacity is better than *TAZ*^*889*^ flies. MLCL can accumulate due to deacylation of cardiolipin during the remodeling processes or due to cardiolipin’s degradation. The improved MLCL:CL ratio in *TAZ*^*889*^ flies overexpressing *spargel* is likely due to decreased CL degradation, since total MLCL did not change, but the total CL did increase. This hypothesis is supported by previous results which also found that the MLCL:CL ratio improved in the flight muscles of flies overexpressing *spargel* (Xu et al., 2019). CL degradation is slowed when CL is associated with other membrane proteins (Xu et al., 2016, Xu et al., 2019). The degradation of CL is likely delayed in *TAZ*^*889*^ flies overexpressing *spargel* due to an increase in mitochondrial membrane proteins (Kaufman et al., 2007, Ventura-Clapier et al., 2008).

In the context of a mitochondrial disease like Barth syndrome, it is reasonable to be cautious about increasing the number of poorly functioning mitochondria, with the possibility of increasing oxidative damage. However, upregulation of *pgc-1α* also increases various antioxidant defense systems to neutralize oxidative species produced from increased oxidative phosphorylation (Austin and St-Pierre, 2012). The ratio between GSH and GSSG, which is a marker of oxidative stress (Kadiiska et al., 2000, Singh et al., 2012), was not significantly higher in *TAZ*^*889*^ flies when they overexpressed *spargel* suggesting that if there is an elevated amount of oxidative stress, the amount is not enough to overwhelm the antioxidant mechanisms to cause substantial damage.

Muscle weakness is a diagnostic characteristic of Barth syndrome (Bittel et al., 2018), so reduced exercise capacity in a muscle-specific *tafazzin* knockdown fly was expected. The improvement to endurance observed when *TAZ* is restored specifically in neurons is intriguing. Little is known about the role neurons play in the pathogenesis of Barth syndrome, but cognitive dysfunction is associated with Barth patients (Mazzocco and Kelley, 2001, Mazzocco et al., 2007). Aging studies demonstrate a correlation between an altered CL profile due to oxidative damage and poor neuronal function (Sen et al., 2007, Lesnefsky and Hoppel, 2008, Petrosillo et al., 2008, Falabella et al., 2021). Even though mitochondria in the muscle are still dysfunctional in neuronal-specific *tafazzin* rescue flies, the endurance and climbing speed are improved by age day 12. Thus, restoration of *tafazzin* in neurons restores baseline exercise capacity without modifying mitochondrial function within muscle tissue. Follow-up studies will be needed to determine what subset of neurons are responsible for this rescue phenotype and how this observation could be leveraged for future therapies.

To our surprise, knocking down *sir2* in *TAZ*^*889*^ flies increased endurance similarly to when *sir2* is overexpressed. However, the mitochondrial function is not restored in *TAZ*^*889*^ flies that have *sir2* knocked down, while the RCR in *sir2* rescue flies is improved. Knockdown of *sir2* in wild-type flies does not increase endurance. Taken together, we believe that knockdown *sir2* in *TAZ* mutants is improving endurance through a different mechanism than overexpressing *sir2*. The increase in endurance of *TAZ*^*889*^ flies with *sir2* knocked down may occur because there is more available NAD^+^ to be utilized by other NAD-dependent enzymes. Although out of the scope of this study, investigation into these enzymes would be valuable as they could be another therapeutic target.

The therapeutic potential of NR, *sir2* and *pgc-1α* for Barth patients is exciting. Other pharmacological stimulators of the NR-Sir2-PGC-1α axis could also have the potential to alleviate symptoms of Barth syndrome. This study demonstrates that NR supplementation provides enormous benefits to the exercise capacity of *tafazzin* mutants, and that it works though *sir2-pgc-1α*. We demonstrated that *sir2* and *pgc-1α* do not need to be overexpressed throughout the whole body for benefits to occur, just in either muscle or neural tissue. Furthermore, overexpression of *sir2* or *pgc-1α* in muscle or neurons can fully mimic restoration of *TAZ*. While gene therapies are still being developed, supplementation with NR is a promising alternative therapy worth further investigation.

## Materials and Methods

### Drosophila lines, Maintenance, and Genetics

Flies were raised at 25°C with 50% humidity. Flies were fed a 10% yeast/sugar diet and kept on a 12-hour light/dark cycle. Fly lines were obtained from the Bloomington *Drosophila* Stock Center (Bloomington, IN), except the following:

- *TAZ*^*889*^ (WellGenetics Inc.)
- Δ*TAZ*/Cyo (Mindong Ren, New York University Langone Health)
- *uas-tafazzin/sb*, *ELAVGS-GAL4* (Mindong Ren, New York University Langone Health)
- *uas-srl* (David Walker, University of California, Las Angeles)

Standard cross schemes were used to create the following lines:

- *TAZ*^*889*^; *MHC-GS-Gal4*
- *TAZ*^*889*^; *ELAV-GS-Gal4*
- *TAZ*^*889*^; *TUB5-GS-Gal4*
- *TAZ*^*889*^; *uas-srl*
- *TAZ*^*889*^; *srl*^*1*^
- *TAZ*^*889*^; *uas-sir2* RNAi,
- *TAZ*^*889*^-*uas-sir2*

The *TAZ*^*2*^, which is shown in Fig. 5 and supplementary Fig. 3, is a trans-heterozygous *tafazzin* mutant with one *TAZ*^*889*^ allele and one Δ*TAZ* allele (Xu et al., 2006).

“GS” denotes a gene-switch line which expresses the Gal4 protein when fed mifepristone (100 μM dissolved in 70% Ethanol). Flies fed mifepristone are represented with “On”, while flies fed the vehicle are represented with “Off.” Using this method controls for genetic background effects since the flies are isogenic, differing only in whether they received the inducing drug. Additionally, this method allows for all gene expression experiments to occur after adult eclosion to avoid any developmental phenotypes. Experimental flies collected within a 48-hour period after eclosing were considered age-matched. Flies were fed mifepristone at age day 2 and were assessed at the earliest on day 5, allowing for at least two full days for induced gene expression changes to accumulate.

### Creation of the *TAZ*^*889*^ mutant using CRSPR/Cas9 genome editing

*TAZ*^*889*^ flies were created (WellGenetics Inc., Taiwan) using CRISPR/Cas-9-mediated genomic editing by homology dependent repair. The 8^th^-896^th^ nucleotides of the *tafazzin* gene were deleted since this size deletion produced a prominent phenotype in a previously published *tafazzin* mutant (Xu et al., 2006). In place of the deletion, a red fluorescent protein (RFP) marker was knocked in to track the presence of the mutation through various cross schemes. All new transgenic experimental flies were “gene-switch” lines and contained the were compared to the identical hybrid background without the inducing drug. The CRISPR plasmid containing the deletion was injected into a *w*^*1118*^ line. That progenitor line was assessed next to *TAZ*^*889*^ flies and served as a genetic background control. The line was validated by PCR and genomic sequencing. Sequencing and Blast results confirmed the deletion and RFP insertion.

### Drug Treatment

Nicotinamide riboside (TRU NIAGEN) was dissolved in water to create 1mM concentration, unless stated otherwise. 50 μl of the solution was applied to the surface of the food and allowed to dry. Flies were placed onto the food at age day 4 and allowed 5 full days on the drug food before assessments.

### Two-phase extraction metabolomics and lipidomics analyses

Metabolomics and lipidomics analyses were performed in 2-5 mg of freeze-dried flies following a two-phase extraction method (Molenaars et al., 2021). Samples were homogenized using a TissueLyser II device (Qiagen; 5 min at 30 pulses/second) in 425 μL water, 500 μL methanol, and 175 μL internal standards mixture. After homogenization, 1000 μL chloroform were added and samples were thoroughly mixed and centrifuged (5 min, 16000 x g, 4°C), creating a two-phase system. The top polar phase was transferred to clean tubes and dried in a vacuum evaporator at 60°C. The bottom apolar fraction was transferred to glass vials and evaporated under a stream of nitrogen at 45 °C.

For metabolomics analysis, pellets obtained after evaporation of the polar phase were dissolved in 100 μL methanol/water (6/4; v/v) and analyzed in an Aquity UPLC system (Waters), coupled to an Impact II Ultra-High Resolution Qq-Time-Of-Flight MS (Bruker). Metabolites were chromatographically separated in a SeQuant ZIC-cHILIC column (PEEK 100 x 2.1 mm, 3 μm particle size; Merck) at 30°C using a method consisting of a gradient running at 0.25 mL/min from 100% mobile phase B (9:1 acetonitrile:water containing 5 mM ammonium acetate pH 6.8) to 100% mobile phase A (1:9 acetonitrile:water containing 5 mM ammonium acetate pH 6.8) for 28 minutes, followed by a re-equilibration step at 100% B for 5 minutes. MS data were acquired in negative and positive ionization modes over the range of m/z 50-1200 and analyzed using Bruker TASQ software (Version 2.1.22.1 1065).

For lipidomics analysis, after evaporation of the apolar fraction, pellets were dissolved in 150 μL chloroform/methanol (1/1; v/v) and analyzed in a Thermo Fisher Scientific Ultimate 3000 binary UPLC coupled to a Q Exactive Plus Orbitrap mass spectrometer using nitrogen as a nebulizing gas. The spray voltage was set at 2500 V, and the capillary temperature at 256°C. For normal-phase separation, a Phenomenex® LUNA silica, 250×2 mm, 5 μm 100 Å was used. Mobile phase consisted of (A) 85:15 (v/v) methanol:water containing 0.0125% formic acid and 3.35 mM ammonia and (B) 97:3 (v/v) chloroform:methanol containing 0.0125% formic acid. The LC gradient consisted of 10% A for 0-1 min, reach 20% A at 4 min, reach 85% A at 12 min, reach 100% A at 12.1 min, 100% A for 12.1-14 min, reach 10% A at 14.1 min, 10% A for 14.1-15 min at a flow rate of 0.3 mL/min and a column temperature of 25 °C. For reverse-phase separation, a Waters HSS T3 column, 150×2.1 mm, 1.8 μm, was used. Mobile phase consisted of (A) 4:6 (v/v) methanol:water and B 1:9 (v/v) methanol:isopropanol, containing 0.1% formic acid and 10 mM ammonia in both cases. The LC gradient consisted of 100% A at 0 min, reach 80% A at 1 min, reach 0% A at 16 min, 0% A for 16-20 min, reach 100% A at 20.1 min, 100% A for 20.1-21 min at a flow rate of 0.4 mL/min and a column temperature of 60 °C. MS data were acquired using negative and positive ionization using continuous scanning over the range of m/z 150-2000. Data were analyzed using an in-house developed metabolomics pipeline written in the R programming language (Molenaars et al., 2021).

All reported metabolite and lipid abundances were normalized to total dry weight as well as to internal standards with comparable retention times and response in the MS.

### Endurance

The endurance of flies was measured as previously described (Tinkerhess et al., 2012a, Damschroder et al., 2018). Eight vials (n = 20 flies) from each cohort were placed onto the Power Tower machine, which stimulates flies to run upwards within their vial until they are physically fatigued. When 80% of the flies within a vial have stopped climbing a minimum of one body length upwards, the vial was removed, and the time was recorded. Survival curves were created in GraphPad Prism (San Diego, CA, USA) and significance was determined using log-rank tests. All assessments were repeated in either duplicate or triplicate. All endurance graphs in the figures show complete data from one repetition and show the same statistical differences as the other replicates unless otherwise stated.

### Climbing Speed

Climbing speed was measured by performing the Rapid Negative Geotaxis (RING) assessment (Gargano et al., 2005). Flies were moved to a clear polypropylene vial (n=100 flies total, 20 flies per vial, 5 vials), and allowed to acclimate for one minute. Vials were moved to a RING apparatus, which allows for five vials to be moved at once. Flies were tapped down to the bottom of their vials and allowed two seconds to climb upwards. After two seconds, a picture was taken to capture their climbing height. Four pictures were taken per group.

ImageJ (Bethesda, MD) was used to quantify the height climbed in two seconds, expressed in quadrants. The average of the four pictures was calculated and plotted. Significance was determined using a Students *t*-test. The climbing speed of at least two separate cohorts was assessed, with 100 flies per cohort. One representative repetition is displayed.

### Mitochondrial Isolation

Isolated mitochondria from 60 flies were used for each biological replicate. Flies were anesthetized on ice for one minute. When heads and thoraces were analyzed, 70-80 fly heads and thoraces were collected. Carcasses were placed in 500 μl of isolation buffer (.32 M sucrose, 10 mM EDTA, 10 mM Tris HCL, 2% fatty-acid free BSA, pH 7.3) (Ferguson et al., 2005). Flies were homogenized using a glass-Teflon Dounce homogenizer (Ferguson et al., 2005, Holmbeck et al., 2015). The homogenate was then filtered through a nylon filter (sigma, pore size = 10 uM) (Ferguson et al., 2005). The filter was washed with 1 mL of isolation buffer. The homogenate was centrifuged at 300 *g* for 5 minutes at 4°C (Holmbeck et al., 2015). The supernatant was moved to a new tube and centrifuged at 6000 *g* for 10 minutes at 4°C. The mitochondrial enriched pellet was resuspended in 30-50 μl respiration buffer (120 mM KCL, 5 mM KH2PO4, 3 mM HEPES, 1 mM EGTA, pH 7.2, BSA free) (Ferguson et al., 2005). Protein concentration was determined using the BCA assay (Thermo Scientific, Rockford, IL, USA). Mitochondrial respiration of isolated mitochondria was measured within two hours of isolation.

### Mitochondrial Respiration

Respiration rates were obtained using a Clark-type electrode (Hansatech Instruments, Norfolk, UK). For all experiments, the chamber temperature was 25°C and 10 μl of isolated mitochondria were added to 990 ml of respiration buffer supplemented with 0.3% BSA. The substrates pyruvate and malate were added to the chamber for a final concentration for 10 uM. State III respiration was induced by addition of ADP [125 nmol] and state IV was induced by adding oligomycin [2.5 um]. The respiratory control ratio was calculated from the respiration rates of state III to state IV. Two technical repetitions were performed for each biological replicate. In total, 6 biological replicates were analyzed and graphed. Statistical significance was determined using either a Students *t*-test or a 2-way ANOVA with *post-hoc* Tukey multiple comparison tests when appropriate. A p-value of less than 0.05 was considered significant.

### qRT-PCR

Gene expression was confirmed by qRT-PCR. Three independent biological replicates were examined with three technical repetitions performed for each. To confirm the RNAi efficacy in the heart specific and fat body specific *tafazzin* knockdown, cDNA was isolated from 20 adult fly hearts and 10 adult fly fat bodies (Sujkowski et al., 2020), using a Cells to CT Kit (Invitrogen, Waltham, MA, USA). Relative abundance of *tafazzin* was determined by amplification and staining with SYBR GREEN and measuring florescence using StudioQuant 3 Real-time PCR System (ThermoFisher, Waltham, MA, USA). For all other genotypes, total RNA or RNA from either the heads or thoraxes of flies, was isolated using TRIZOL (Invitrogen, Waltham, MA, USA). One step qRT-PCR was performed using *Power* SYBR Green master mix (Applied Biosystems, Waltham, MA, USA) and was performing using StudioQuant 3 Real-time PCR System (ThermoFisher, Waltham, MA, USA). Each 20 μl reaction contained of RNA (4 μl, 25 ng/μl), forward and reverse primers (2 μl, 1 μM), *Power SYBR Green PCR Master Mix* (10 μl) reverse transcriptase (0.1 μl), inhibitor (0.025 μl), and dH2O (3.875 μl). The qPCR program ran for 40 cycles of 95 °C for 15 seconds followed by 60 °C for 1 minute. mRNA data were normalized to *Act5C*.

List of Primers:

Act5C F: CGCAGAGCAAGCGTGGTA
Act5C R: GTGCCACACGCAGCTCAT
*tafazzin* F: CGTGTGTTCCAATTTAAGGCGT
tafazzin R: AGTCTGGGTGGCGATATCCT
*srl* F: GGATTCACGAATGCTAAATGTGTTCC
*srl* R: GATGGGTAGGATGCCGCTCAG
*sir2* F: GCCTCCAGGACAGTTAGCAG
*sir2* RGCCAATCTCTTGTTCTTCCC

### mtDNA Copy Number

DNA was isolated from 20 flies using phenol-chlorofrm (Sigma Aldrich, St. Louis, MO, USA) (n = 20 flies per replicate group). mtDNA was quantified by amplifying the mitochondrial large ribosomal RNA gene (lrRNA), which does not have a nuclear copy in *Drosophila melanogaster* (Correa et al., 2012), and the nuclear genomic DNA was quantified by amplifying RNA polymerase II (Correa et al., 2012).

Quantitative PCR (qPCR) (Aw et al., 2018) was performed using a ABI 7300 Real Time PCR System (Applied Biosystems, Waltham, MA, USA) and *Power SYBR Green PCR Master Mix* (Applied Biosystems, Waltham, MA, USA). The qPCR program started with a denaturing step at 95 °C for 5 minutes followed by amplification for 40 cycles that consisted of 95 °C for 5 minutes followed by 60 °C for 30 seconds (Aw et al., 2018). The mtDNA copy number is expressed as the average fold change of mtDNA to nuclear DNA (Aw et al., 2018). Primers were previously published (Correa et al., 2012). The data was analyzed using either a student’s *t-test*, or a 2-way ANOVA with *post-hoc* Tukey multiple comparison tests.

Primer Sequences:

lrRNA 5’: TCGTCCAACCATTCATTCC
lrRNA 3’: ATAAAGTCTAACCTGCCCACTGA
rp2 5’: AGGCGTTTGAGTGGTTGG
rp2 3’: TGGAAGGTGTTCAGTGTCAT

## Acknowledgements

We thank Maik Huttemann for his guidance with the mitochondrial isolation protocol and Clark electrode measurements. We would also like to thank Miriam Greenberg, Jiajia Ji, and Christian Reynolds for their helpful advice. We thank David Walker and Mindong Ren for sending fly lines.

## Author contributions statement

DD designed the experiments, analyzed data, and wrote the manuscript. RZP designed experiments regarding mass spectrometry and analyzed the data. RW and RHH designed experiments, analyzed data, edited manuscript, and provided funding. All authors have read and approved the final version of the manuscript.

## References

Amjad, S., Nisar, S., Bhat, A. A., Shah, A. R., Frenneaux, M. P., Fakhro, K., Haris, M., Reddy, R., Patay, Z., Baur, J. & Bagga, P. 2021. Role of NAD+ in regulating cellular and metabolic signaling pathways. Molecular Metabolism, 49, 101195.

Austin, S. & St-Pierre, J. 2012. PGC1alpha and mitochondrial metabolism--emerging concepts and relevance in ageing and neurodegenerative disorders. J Cell Sci, 125, 4963–71.

Aw, W. C., Towarnicki, S. G., Melvin, R. G., Youngson, N. A., Garvin, M. R., Hu, Y., Nielsen, S., Thomas, T., Pickford, R., Bustamante, S., Vila-Sanjurjo, A., Smyth, G. K. & Ballard, J. W. O. 2018. Genotype to phenotype: Diet-by-mitochondrial DNA haplotype interactions drive metabolic flexibility and organismal fitness. PLoS Genet, 14, e1007735.

Baar, K., Wende, A. R., Jones, T. E., Marison, M., Nolte, L. A., Chen, M., Kelly, D. P. & Holloszy, J. O. 2002. Adaptations of skeletal muscle to exercise: rapid increase in the transcriptional coactivator PGC-1. FASEB J, 16, 1879–86.

Barth, P. G., Scholte, H. R., Berden, J. A., Van Der Klei-Van Moorsel, J. M., Luyt-Houwen, I. E., Van ’T Veer-Korthof, E. T., Van Der Harten, J. J. & Sobotka-Plojhar, M. A. 1983. An X-linked mitochondrial disease affecting cardiac muscle, skeletal muscle and neutrophil leucocytes. J Neurol Sci, 62, 327–55.

Berrisford, J. M. & Sazanov, L. A. 2009. Structural basis for the mechanism of respiratory complex I. J Biol Chem, 284, 29773–83.

Bittel, A. J., Bohnert, K. L., Reeds, D. N., Peterson, L. R., De Las Fuentes, L., Corti, M., Taylor, C. L., Byrne, B. J. & Cade, W. T. 2018. Reduced Muscle Strength in Barth Syndrome May Be Improved by Resistance Exercise Training: A Pilot Study. JIMD Rep, 41, 63–72.

Bowen, V., Milligan, E., Mccormack, S., Mccurdy, M., Smart, T., Sheriff, L., Toth, M. & Valentine, J. 2019. The Voice of the Patient: Barth Syndrome.

Bowron, A., Honeychurch, J., Williams, M., Tsai-Goodman, B., Clayton, N., Jones, L., Shortland, G. J., Qureshi, S. A., Heales, S. J. & Steward, C. G. 2015. Barth syndrome without tetralinoleoyl cardiolipin deficiency: a possible ameliorated phenotype. J Inherit Metab Dis, 38, 279–86.

Brown, K. D., Maqsood, S., Huang, J. Y., Pan, Y., Harkcom, W., Li, W., Sauve, A., Verdin, E. & Jaffrey, S. R. 2014. Activation of SIRT3 by the NAD^+^ precursor nicotinamide riboside protects from noise-induced hearing loss. Cell Metab, 20, 1059–68.

Cade, W. T., Reeds, D. N., Peterson, L. R., Bohnert, K. L., Tinius, R. A., Benni, P. B., Byrne, B. J. & Taylor, C. L. 2017. Endurance Exercise Training in Young Adults with Barth Syndrome: A Pilot Study. JIMD Rep, 32, 15–24.

Cambronne, X. A. & Kraus, W. L. 2020. Location, Location, Location: Compartmentalization of NAD+ Synthesis and Functions in Mammalian Cells. Trends in Biochemical Sciences, 45, 858–873.

Canto, C., Houtkooper, R. H., Pirinen, E., Youn, D. Y., Oosterveer, M. H., Cen, Y., Fernandez-Marcos, P. J., Yamamoto, H., Andreux, P. A., Cettour-Rose, P., Gademann, K., Rinsch, C., Schoonjans, K., Sauve, A. A. & Auwerx, J. 2012. The NAD(+) precursor nicotinamide riboside enhances oxidative metabolism and protects against high-fat diet-induced obesity. Cell Metab, 15, 838–47.

Cerutti, R., Pirinen, E., Lamperti, C., Marchet, S., Sauve, A. A., Li, W., Leoni, V., Schon, E. A., Dantzer, F., Auwerx, J., Viscomi, C. & Zeviani, M. 2014. NAD(+)-dependent activation of Sirt1 corrects the phenotype in a mouse model of mitochondrial disease. Cell Metab, 19, 1042–9.

Clement, J., Wong, M., Poljak, A., Sachdev, P. & Braidy, N. 2019. The Plasma NAD(+) Metabolome Is Dysregulated in “Normal” Aging. Rejuvenation Res, 22, 121–130.

Conze, D., Brenner, C. & Kruger, C. L. 2019. Safety and Metabolism of Long-term Administration of NIAGEN (Nicotinamide Riboside Chloride) in a Randomized, Double-Blind, Placebo-controlled Clinical Trial of Healthy Overweight Adults. Sci Rep, 9, 9772.

Conze, D. B., Crespo-Barreto, J. & Kruger, C. L. 2016. Safety assessment of nicotinamide riboside, a form of vitamin B3. Hum Exp Toxicol, 35, 1149–1160.

Correa, C. C., Aw, W. C., Melvin, R. G., Pichaud, N. & Ballard, J. W. 2012. Mitochondrial DNA variants influence mitochondrial bioenergetics in Drosophila melanogaster. Mitochondrion, 12, 459–64.

Damschroder, D., Reynolds, C. & Wessells, R. 2018. Drosophila tafazzin mutants have impaired exercise capacity. Physiol Rep, 6.

Deena Damschroder, T. C., Alyson Sujkowski, Robert Wessells 2018. *Drosophila* Endurance Training and Assessment of Its Effects on Systemic Adaptions. BioProtocol, 8.

Elhassan, Y. S., Kluckova, K., Fletcher, R. S., Schmidt, M. S., Garten, A., Doig, C. L., Cartwright, D. M., Oakey, L., Burley, C. V., Jenkinson, N., Wilson, M., Lucas, S. J. E., Akerman, I., Seabright, A., Lai, Y. C., Tennant, D. A., Nightingale, P., Wallis, G. A., Manolopoulos, K. N., Brenner, C., Philp, A. & Lavery, G. G. 2019. Nicotinamide Riboside Augments the Aged Human Skeletal Muscle NAD(+) Metabolome and Induces Transcriptomic and Anti-inflammatory Signatures. Cell Rep, 28, 1717–1728 e6.

Falabella, M., Vernon, H. J., Hanna, M. G., Claypool, S. M. & Pitceathly, R. D. S. 2021. Cardiolipin, Mitochondria, and Neurological Disease. Trends Endocrinol Metab, 32, 224–237.

Fang, E. F., Hou, Y., Lautrup, S., Jensen, M. B., Yang, B., Sengupta, T., Caponio, D., Khezri, R., Demarest, T. G., Aman, Y., Figueroa, D., Morevati, M., Lee, H. J., Kato, H., Kassahun, H., Lee, J. H., Filippelli, D., Okur, M. N., Mangerich, A., Croteau, D. L., Maezawa, Y., Lyssiotis, C. A., Tao, J., Yokote, K., Rusten, T. E., Mattson, M. P., Jasper, H., Nilsen, H. & Bohr, V. A. 2019. NAD(+) augmentation restores mitophagy and limits accelerated aging in Werner syndrome. Nat Commun, 10, 5284.

Fang, E. F., Lautrup, S., Hou, Y., Demarest, T. G., Croteau, D. L., Mattson, M. P. & Bohr, V. A. 2017. NAD(+) in Aging: Molecular Mechanisms and Translational Implications. Trends Mol Med, 23, 899–916.

Fang, E. F., Scheibye-Knudsen, M., Chua, K. F., Mattson, M. P., Croteau, D. L. & Bohr, V. A. 2016. Nuclear DNA damage signalling to mitochondria in ageing. Nat Rev Mol Cell Biol, 17, 308–21.

Ferguson, M., Mockett, R. J., Shen, Y., Orr, W. C. & Sohal, R. S. 2005. Age-associated decline in mitochondrial respiration and electron transport in Drosophila melanogaster. Biochem J, 390, 501–11.

Fry, M. & Green, D. E. 1980. Cardiolipin requirement by cytochrome oxidase and the catalytic role of phospholipid. Biochem Biophys Res Commun, 93, 1238–46.

Garavaglia, S., Bruzzone, S., Cassani, C., Canella, L., Allegrone, G., Sturla, L., Mannino, E., Millo, E., De Flora, A. & Rizzi, M. 2012. The high-resolution crystal structure of periplasmic Haemophilus influenzae NAD nucleotidase reveals a novel enzymatic function of human CD73 related to NAD metabolism. Biochem J, 441, 131–41.

Gargano, J. W., Martin, I., Bhandari, P. & Grotewiel, M. S. 2005. Rapid iterative negative geotaxis (RING): a new method for assessing age-related locomotor decline in Drosophila. Exp Gerontol, 40, 386–95.

Gonzalvez, F., D’aurelio, M., Boutant, M., Moustapha, A., Puech, J. P., Landes, T., Arnaune-Pelloquin, L., Vial, G., Taleux, N., Slomianny, C., Wanders, R. J., Houtkooper, R. H., Bellenguer, P., Moller, I. M., Gottlieb, E., Vaz, F. M., Manfredi, G. & Petit, P. X. 2013. Barth syndrome: cellular compensation of mitochondrial dysfunction and apoptosis inhibition due to changes in cardiolipin remodeling linked to tafazzin (TAZ) gene mutation. Biochim Biophys Acta, 1832, 1194–206.

Grozio, A., Mills, K. F., Yoshino, J., Bruzzone, S., Sociali, G., Tokizane, K., Lei, H. C., Cunningham, R., Sasaki, Y., Migaud, M. E. & Imai, S. I. 2019. Slc12a8 is a nicotinamide mononucleotide transporter. Nat Metab, 1, 47–57.

Grozio, A., Sociali, G., Sturla, L., Caffa, I., Soncini, D., Salis, A., Raffaelli, N., De Flora, A., Nencioni, A. & Bruzzone, S. 2013. CD73 protein as a source of extracellular precursors for sustained NAD+ biosynthesis in FK866-treated tumor cells. J Biol Chem, 288, 25938–25949.

Handschin, C., Chin, S., Li, P., Liu, F., Maratos-Flier, E., Lebrasseur, N. K., Yan, Z. & Spiegelman, B. M. 2007. Skeletal muscle fiber-type switching, exercise intolerance, and myopathy in PGC-1alpha muscle-specific knock-out animals. J Biol Chem, 282, 30014–21.

Holmbeck, M. A., Donner, J. R., Villa-Cuesta, E. & Rand, D. M. 2015. A Drosophila model for mito-nuclear diseases generated by an incompatible interaction between tRNA and tRNA synthetase. Dis Model Mech, 8, 843–54.

Hostetler, K. Y., Van Den Bosch, H. & Van Deenen, L. L. 1971. Biosynthesis of cardiolipin in liver mitochondria. Biochim Biophys Acta, 239, 113–9.

Houtkooper, R. H., Rodenburg, R. J., Thiels, C., Van Lenthe, H., Stet, F., Poll-The, B. T., Stone, J. E., Steward, C. G., Wanders, R. J., Smeitink, J., Kulik, W. & Vaz, F. M. 2009a. Cardiolipin and monolysocardiolipin analysis in fibroblasts, lymphocytes, and tissues using high-performance liquid chromatography-mass spectrometry as a diagnostic test for Barth syndrome. Anal Biochem, 387, 230–7.

Houtkooper, R. H., Turkenburg, M., Poll-The, B. T., Karall, D., Pérez-Cerdá, C., Morrone, A., Malvagia, S., Wanders, R. J., Kulik, W. & Vaz, F. M. 2009b. The enigmatic role of tafazzin in cardiolipin metabolism. Biochimica et Biophysica Acta (BBA) - Biomembranes, 1788, 2003–2014.

Imai, S. I. & Guarente, L. 2016. It takes two to tango: NAD(+) and sirtuins in aging/longevity control. NPJ Aging Mech Dis, 2, 16017.

Kadiiska, M. B., Gladen, B. C., Baird, D. D., Dikalova, A. E., Sohal, R. S., Hatch, G. E., Jones, D. P., Mason, R. P. & Barrett, J. C. 2000. Biomarkers of oxidative stress study: are plasma antioxidants markers of CCl(4) poisoning? Free Radic Biol Med, 28, 838–45.

Kamanna, V. S., Ganji, S. H. & Kashyap, M. L. 2009. The mechanism and mitigation of niacin-induced flushing. Int J Clin Pract, 63, 1369–77.

Kaufman, B. A., Durisic, N., Mativetsky, J. M., Costantino, S., Hancock, M. A., Grutter, P. & Shoubridge, E. A. 2007. The mitochondrial transcription factor TFAM coordinates the assembly of multiple DNA molecules into nucleoid-like structures. Mol Biol Cell, 18, 3225–36.

Kelley, R. I., Cheatham, J. P., Clark, B. J., Nigro, M. A., Powell, B. R., Sherwood, G. W., Sladky, J. T. & Swisher, W. P. 1991. X-linked dilated cardiomyopathy with neutropenia, growth retardation, and 3-methylglutaconic aciduria. J Pediatr, 119, 738–47.

Khan, N. A., Auranen, M., Paetau, I., Pirinen, E., Euro, L., Forsstrom, S., Pasila, L., Velagapudi, V., Carroll, C. J., Auwerx, J. & Suomalainen, A. 2014. Effective treatment of mitochondrial myopathy by nicotinamide riboside, a vitamin B3. EMBO Mol Med, 6, 721–31.

Kulik, W., Van Lenthe, H., Stet, F. S., Houtkooper, R. H., Kemp, H., Stone, J. E., Steward, C. G., Wanders, R. J. & Vaz, F. M. 2008. Bloodspot assay using HPLC-tandem mass spectrometry for detection of Barth syndrome. Clin Chem, 54, 371–8.

Lai, L., Wang, M., Martin, O. J., Leone, T. C., Vega, R. B., Han, X. & Kelly, D. P. 2014. A role for peroxisome proliferator-activated receptor gamma coactivator 1 (PGC-1) in the regulation of cardiac mitochondrial phospholipid biosynthesis. J Biol Chem, 289, 2250–9.

Lautrup, S., Sinclair, D. A., Mattson, M. P. & Fang, E. F. 2019. NAD(+) in Brain Aging and Neurodegenerative Disorders. Cell Metab, 30, 630–655.

Lehman, J. J., Barger, P. M., Kovacs, A., Saffitz, J. E., Medeiros, D. M. & Kelly, D. P. 2000. Peroxisome proliferator-activated receptor gamma coactivator-1 promotes cardiac mitochondrial biogenesis. J Clin Invest, 106, 847–56.

Lehmann, S., Loh, S. H. & Martins, L. M. 2017. Enhancing NAD(+) salvage metabolism is neuroprotective in a PINK1 model of Parkinson’s disease. Biol Open, 6, 141–147.

Lesnefsky, E. J. & Hoppel, C. L. 2008. Cardiolipin as an oxidative target in cardiac mitochondria in the aged rat. Biochimica Et Biophysica Acta-Bioenergetics, 1777, 1020–1027.

Li, Y., Xu, W., Mcburney, M. W. & Longo, V. D. 2008. Sirt1 inhibition reduces IGF-I/IRS-2/Ras/ERK1/2 signaling and protects neurons. Cell Metab, 8, 38–48.

Liu, L., Su, X., Quinn, W. J., 3rd, Hui, S., Krukenberg, K., Frederick, D. W., Redpath, P., Zhan, L., Chellappa, K., White, E., Migaud, M., Mitchison, T. J., Baur, J. A. & Rabinowitz, J. D. 2018. Quantitative Analysis of NAD Synthesis-Breakdown Fluxes. Cell Metab, 27, 1067–1080 e5.

Martens, C. R., Denman, B. A., Mazzo, M. R., Armstrong, M. L., Reisdorph, N., Mcqueen, M. B., Chonchol, M. & Seals, D. R. 2018. Chronic nicotinamide riboside supplementation is well-tolerated and elevates NAD(+) in healthy middle-aged and older adults. Nat Commun, 9, 1286.

Massudi, H., Grant, R., Braidy, N., Guest, J., Farnsworth, B. & Guillemin, G. J. 2012. Age-associated changes in oxidative stress and NAD+ metabolism in human tissue. PLoS One, 7, e42357.

Mazzocco, M. M., Henry, A. E. & Kelly, R. I. 2007. Barth syndrome is associated with a cognitive phenotype. J Dev Behav Pediatr, 28, 22–30.

Mazzocco, M. M. & Kelley, R. I. 2001. Preliminary evidence for a cognitive phenotype in Barth syndrome. Am J Med Genet, 102, 372–8.

Mckenzie, M., Lazarou, M., Thorburn, D. R. & Ryan, M. T. 2006. Mitochondrial respiratory chain supercomplexes are destabilized in Barth Syndrome patients. J Mol Biol, 361, 462–9.

Mehmel, M., Jovanović, N. & Spitz, U. 2020. Nicotinamide Riboside-The Current State of Research and Therapeutic Uses. Nutrients, 12.

Michishita, E., Park, J. Y., Burneskis, J. M., Barrett, J. C. & Horikawa, I. 2005. Evolutionarily conserved and nonconserved cellular localizations and functions of human SIRT proteins. Mol Biol Cell, 16, 4623–35.

Molenaars, M., Schomakers, B. V., Elfrink, H. L., Gao, A. W., Vervaart, M. a. T., Pras-Raves, M. L., Luyf, A. C., Smith, R. L., Sterken, M. G., Kammenga, J. E., Van Kampen, A. H. C., Janssens, G. E., Vaz, F. M., Van Weeghel, M. & Houtkooper, R. H. 2021. Metabolomics and lipidomics in Caenorhabditis elegans using a single-sample preparation. Dis Model Mech, 14.

Mukherjee, S., Basar, M. A., Davis, C. & Duttaroy, A. 2014. Emerging functional similarities and divergences between Drosophila Spargel/dPGC-1 and mammalian PGC-1 protein. Front Genet, 5, 216.

Neustein, H. B., Lurie, P. R., Dahms, B. & Takahashi, M. 1979. An X-linked recessive cardiomyopathy with abnormal mitochondria. Pediatrics, 64, 24–9.

Neuwald, A. F. 1997. Barth syndrome may be due to an acyltransferase deficiency. Curr Biol, 7, R465–6.

Olesen, J., Kiilerich, K. & Pilegaard, H. 2010. PGC-1alpha-mediated adaptations in skeletal muscle. Pflugers Arch, 460, 153–62.

Onyango, P., Celic, I., Mccaffery, J. M., Boeke, J. D. & Feinberg, A. P. 2002. SIRT3, a human SIR2 homologue, is an NAD-dependent deacetylase localized to mitochondria. Proc Natl Acad Sci U S A, 99, 13653–8.

Petrosillo, G., Matera, M., Casanova, G., Ruggiero, F. M. & Paradies, G. 2008. Mitochondrial dysfunction in rat brain with aging Involvement of complex I, reactive oxygen species and cardiolipin. Neurochemistry International, 53, 126–131.

Pfeiffer, K., Gohil, V., Stuart, R. A., Hunte, C., Brandt, U., Greenberg, M. L. & Schagger, H. 2003. Cardiolipin stabilizes respiratory chain supercomplexes. J Biol Chem, 278, 52873–80.

Pirinen, E., Auranen, M., Khan, N. A., Brilhante, V., Urho, N., Pessia, A., Hakkarainen, A., Kuula, J., Heinonen, U., Schmidt, M. S., Haimilahti, K., Piirila, P., Lundbom, N., Taskinen, M. R., Brenner, C., Velagapudi, V., Pietilainen, K. H. & Suomalainen, A. 2020. Niacin Cures Systemic NAD(+) Deficiency and Improves Muscle Performance in Adult-Onset Mitochondrial Myopathy. Cell Metab, 31, 1078–1090 e5.

Rajman, L., Chwalek, K. & Sinclair, D. A. 2018. Therapeutic Potential of NAD-Boosting Molecules: The In Vivo Evidence. Cell Metab, 27, 529–547.

Ratajczak, J., Joffraud, M., Trammell, S. A., Ras, R., Canela, N., Boutant, M., Kulkarni, S. S., Rodrigues, M., Redpath, P., Migaud, M. E., Auwerx, J., Yanes, O., Brenner, C. & Canto, C. 2016. NRK1 controls nicotinamide mononucleotide and nicotinamide riboside metabolism in mammalian cells. Nat Commun, 7, 13103.

Remie, C. M. E., Roumans, K. H. M., Moonen, M. P. B., Connell, N. J., Havekes, B., Mevenkamp, J., Lindeboom, L., De Wit, V. H. W., Van De Weijer, T., Aarts, S., Lutgens, E., Schomakers, B. V., Elfrink, H. L., Zapata-Perez, R., Houtkooper, R. H., Auwerx, J., Hoeks, J., Schrauwen-Hinderling, V. B., Phielix, E. & Schrauwen, P. 2020. Nicotinamide riboside supplementation alters body composition and skeletal muscle acetylcarnitine concentrations in healthy obese humans. Am J Clin Nutr, 112, 413–426.

Rera, M., Bahadorani, S., Cho, J., Koehler, C. L., Ulgherait, M., Hur, J. H., Ansari, W. S., Lo, T., Jr., Jones, D. L. & Walker, D. W. 2011. Modulation of longevity and tissue homeostasis by the Drosophila PGC-1 homolog. Cell Metab, 14, 623–34.

Roberts, A. E., Nixon, C., Steward, C. G., Gauvreau, K., Maisenbacher, M., Fletcher, M., Geva, J., Byrne, B. J. & Spencer, C. T. 2012. The Barth Syndrome Registry: distinguishing disease characteristics and growth data from a longitudinal study. Am J Med Genet A, 158A, 2726–32.

Rodgers, J. T., Lerin, C., Haas, W., Gygi, S. P., Spiegelman, B. M. & Puigserver, P. 2005. Nutrient control of glucose homeostasis through a complex of PGC-1alpha and SIRT1. Nature, 434, 113–8.

Sazanov, L. A. 2015. A giant molecular proton pump: structure and mechanism of respiratory complex I. Nat Rev Mol Cell Biol, 16, 375–88.

Schlame, M. 2013. Cardiolipin remodeling and the function of tafazzin. Biochim Biophys Acta, 1831, 582–8.

Schlame, M. & Haldar, D. 1993. Cardiolipin is synthesized on the matrix side of the inner membrane in rat liver mitochondria. J Biol Chem, 268, 74–9.

Schlame, M., Ren, M., Xu, Y., Greenberg, M. L. & Haller, I. 2005. Molecular symmetry in mitochondrial cardiolipins. Chem Phys Lipids, 138, 38–49.

Schlame, M., Towbin, J. A., Heerdt, P. M., Jehle, R., Dimauro, S. & Blanck, T. J. 2002. Deficiency of tetralinoleoyl-cardiolipin in Barth syndrome. Ann Neurol, 51, 634–7.

Schondorf, D. C., Ivanyuk, D., Baden, P., Sanchez-Martinez, A., De Cicco, S., Yu, C., Giunta, I., Schwarz, L. K., Di Napoli, G., Panagiotakopoulou, V., Nestel, S., Keatinge, M., Pruszak, J., Bandmann, O., Heimrich, B., Gasser, T., Whitworth, A. J. & Deleidi, M. 2018. The NAD+ Precursor Nicotinamide Riboside Rescues Mitochondrial Defects and Neuronal Loss in iPSC and Fly Models of Parkinson’s Disease. Cell Rep, 23, 2976–2988.

Sen, T., Sen, N., Jana, S., Khan, F. H., Chatterjee, U. & Chakrabarti, S. 2007. Depolarization and cardiolipin depletion in aged rat brain mitochondria: Relationship with oxidative stress and electron transport chain activity. Neurochemistry International, 50, 719–725.

Singh, S., Khan, A. R. & Gupta, A. K. 2012. Role of glutathione in cancer pathophysiology and therapeutic interventions. J Exp Ther Oncol, 9, 303–16.

Spencer, C. T., Bryant, R. M., Day, J., Gonzalez, I. L., Colan, S. D., Thompson, W. R., Berthy, J., Redfearn, S. P. & Byrne, B. J. 2006. Cardiac and clinical phenotype in Barth syndrome. Pediatrics, 118, e337–46.

Srivastava, S. 2016. Emerging therapeutic roles for NAD(+) metabolism in mitochondrial and age-related disorders. Clinical and translational medicine, 5, 25–25.

Stein, L. R. & Imai, S. 2012. The dynamic regulation of NAD metabolism in mitochondria. Trends Endocrinol Metab, 23, 420–8.

Stocks, B., Ashcroft, S. P., Joanisse, S., Dansereau, L. C., Koay, Y. C., Elhassan, Y. S., Lavery, G. G., Quek, L. E., O’sullivan, J. F., Philp, A. M., Wallis, G. A. & Philp, A. 2021. Nicotinamide riboside supplementation does not alter whole-body or skeletal muscle metabolic responses to a single bout of endurance exercise. J Physiol, 599, 1513–1531.

Sujkowski, A., Gretzinger, A., Soave, N., Todi, S. V. & Wessells, R. 2020. Alpha- and beta-adrenergic octopamine receptors in muscle and heart are required for Drosophila exercise adaptations. PLoS Genet, 16, e1008778.

Suzuki-Hatano, S., Saha, M., Rizzo, S. A., Witko, R. L., Gosiker, B. J., Ramanathan, M., Soustek, M. S., Jones, M. D., Kang, P. B., Byrne, B. J., Cade, W. T. & Pacak, C. A. 2019. AAV-Mediated TAZ Gene Replacement Restores Mitochondrial and Cardioskeletal Function in Barth Syndrome. Hum Gene Ther, 30, 139–154.

Thompson, W. R., Decroes, B., Mcclellan, R., Rubens, J., Vaz, F. M., Kristaponis, K., Avramopoulos, D. & Vernon, H. J. 2016. New targets for monitoring and therapy in Barth syndrome. Genet Med, 18, 1001–10.

Tiefenbock, S. K., Baltzer, C., Egli, N. A. & Frei, C. 2010. The Drosophila PGC-1 homologue Spargel coordinates mitochondrial activity to insulin signalling. EMBO J, 29, 171–83.

Tinkerhess, M. J., Ginzberg, S., Piazza, N. & Wessells, R. J. 2012a. Endurance training protocol and longitudinal performance assays for Drosophila melanogaster. J Vis Exp.

Tinkerhess, M. J., Healy, L., Morgan, M., Sujkowski, A., Matthys, E., Zheng, L. & Wessells, R. J. 2012b. The Drosophila PGC-1alpha homolog spargel modulates the physiological effects of endurance exercise. PLoS One, 7, e31633.

Trammell, S. A., Schmidt, M. S., Weidemann, B. J., Redpath, P., Jaksch, F., Dellinger, R. W., Li, Z., Abel, E. D., Migaud, M. E. & Brenner, C. 2016. Nicotinamide riboside is uniquely and orally bioavailable in mice and humans. Nat Commun, 7, 12948.

Ventura-Clapier, R., Garnier, A. & Veksler, V. 2008. Transcriptional control of mitochondrial biogenesis: the central role of PGC-1α. Cardiovascular Research, 79, 208–217.

Vreken, P., Valianpour, F., Nijtmans, L. G., Grivell, L. A., Plecko, B., Wanders, R. J. & Barth, P. G. 2000. Defective remodeling of cardiolipin and phosphatidylglycerol in Barth syndrome. Biochem Biophys Res Commun, 279, 378–82.

Wang, S., Wan, T., Ye, M., Qiu, Y., Pei, L., Jiang, R., Pang, N., Huang, Y., Liang, B., Ling, W., Lin, X., Zhang, Z. & Yang, L. 2018. Nicotinamide riboside attenuates alcohol induced liver injuries via activation of SirT1/PGC-1alpha/mitochondrial biosynthesis pathway. Redox Biol, 17, 89–98.

Wu, Z., Puigserver, P., Andersson, U., Zhang, C., Adelmant, G., Mootha, V., Troy, A., Cinti, S., Lowell, B., Scarpulla, R. C. & Spiegelman, B. M. 1999. Mechanisms controlling mitochondrial biogenesis and respiration through the thermogenic coactivator PGC-1. Cell, 98, 115–24.

Xu, Y., Anjaneyulu, M., Donelian, A., Yu, W., Greenberg, M. L., Ren, M., Owusu-Ansah, E. & Schlame, M. 2019. Assembly of the complexes of oxidative phosphorylation triggers the remodeling of cardiolipin. Proc Natl Acad Sci U S A, 116, 11235–11240.

Xu, Y., Condell, M., Plesken, H., Edelman-Novemsky, I., Ma, J., Ren, M. & Schlame, M. 2006. A Drosophila model of Barth syndrome. Proc Natl Acad Sci U S A, 103, 11584–8.

Xu, Y., Phoon, C. K., Berno, B., D’souza, K., Hoedt, E., Zhang, G., Neubert, T. A., Epand, R. M., Ren, M. & Schlame, M. 2016. Loss of protein association causes cardiolipin degradation in Barth syndrome. Nat Chem Biol, 12, 641–7.

Yang, Q., Cong, L., Wang, Y., Luo, X., Li, H., Wang, H., Zhu, J., Dai, S., Jin, H., Yao, G., Shi, S., Hsueh, A. J. & Sun, Y. 2020. Increasing ovarian NAD(+) levels improve mitochondrial functions and reverse ovarian aging. Free Radic Biol Med, 156, 1–10.

Yoshino, J., Mills, K. F., Yoon, M. J. & Imai, S. 2011. Nicotinamide mononucleotide, a key NAD(+) intermediate, treats the pathophysiology of diet- and age-induced diabetes in mice. Cell Metab, 14, 528–36.

Yoshino, M., Yoshino, J., Kayser, B. D., Patti, G. J., Franczyk, M. P., Mills, K. F., Sindelar, M., Pietka, T., Patterson, B. W., Imai, S. I. & Klein, S. 2021. Nicotinamide mononucleotide increases muscle insulin sensitivity in prediabetic women. Science, 372, 1224–1229.

Zakhary, S. M., Ayubcha, D., Dileo, J. N., Jose, R., Leheste, J. R., Horowitz, J. M. & Torres, G. 2010. Distribution analysis of deacetylase SIRT1 in rodent and human nervous systems. Anat Rec (Hoboken), 293, 1024–32.

Zapata-Perez, R., Wanders, R. J. A., Van Karnebeek, C. D. M. & Houtkooper, R. H. 2021. NAD(+) homeostasis in human health and disease. EMBO Mol Med, 13, e13943.

Zhou, Q., Zhu, L., Qiu, W., Liu, Y., Yang, F., Chen, W. & Xu, R. 2020. Nicotinamide Riboside Enhances Mitochondrial Proteostasis and Adult Neurogenesis through Activation of Mitochondrial Unfolded Protein Response Signaling in the Brain of ALS SOD1(G93A) Mice. Int J Biol Sci, 16, 284–297.

