## Supplemental figures for "Stimulating the *sir2-pgc-1ɑ* axis rescues exercise capacity and mitochondrial respiration in *Drosophila tafazzin* mutants"

A.

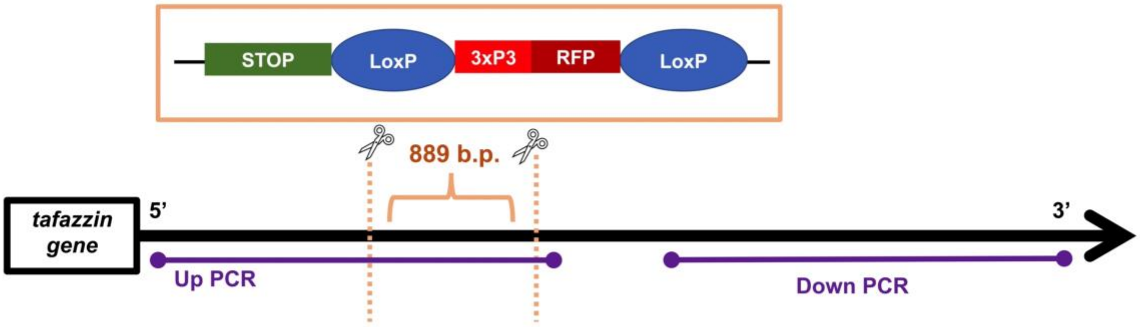


B.

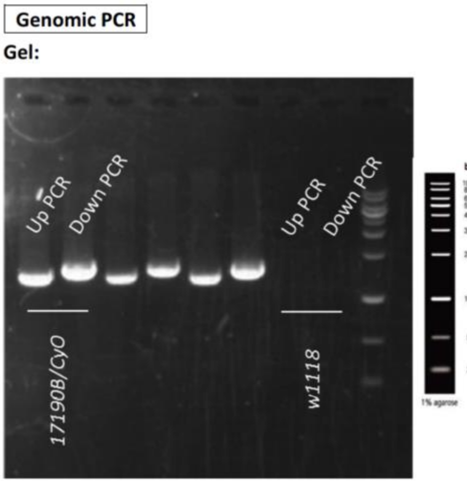


**Supplementary Figure 1: Genotype data for *$TAZ^{889}$* flies.** (A.) The *$TAZ^{889}$* allele was made by deleting 889 base pairs (b.p.) from *tafazzin* using CRISPR Cas-9 mediated genomic editing by homology-dependent repair. In place of the deleted base pairs, a Stop-RFP cassette was inserted, which expresses a red-fluorescent protein in the eyes of *$TAZ^{889}$* flies. The cassette was then injected into *$w^{1118}$* line. Homozygous offspring containing the mutation were then collected and validated using PCR and sequencing. (B.) To confirm the STOP-RFP cassette was inserted into the gene, PCR was performed. The two amplicons are represented by “Up PCR” and “Down PCR”. The forward primer for the UP PCR segment was designed for the 5’ end of the amplicon and the reverse primer was designed for the 3xP3 promoter. Therefore, DNA would only be amplified if the STOP-RFP cassette was inserted into the gene. The Down PCR segment was amplified using a forward primer designed for the alpha-Tubulin 3’ UTR within the STOP-RFP cassette and a reverse primer designed for the 3’ end of the gene. Therefore the Down PCR segment would only be replicated if the STOP-RFP cassette was inserted into the gene. PCR bands for both the Up (1368 b.p.) and Down PCR (1540 b.p.) segments were of expected sizes. No bands were observed in the control lines (*$w^{1118}$*). The PCR products were then sent for sequencing, which confirmed the 889 base pair deletion. The creation and validation of the *$TAZ^{889}$* allele was performed by WellGenetics Inc.

**A.**

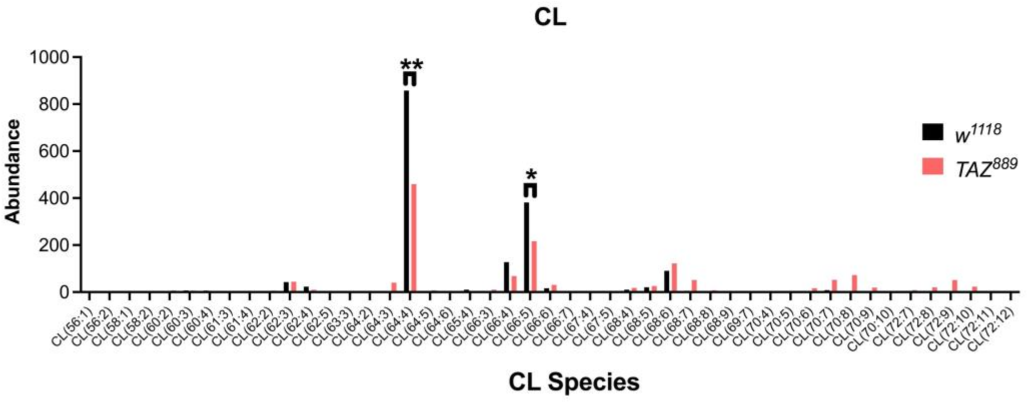


**B.**

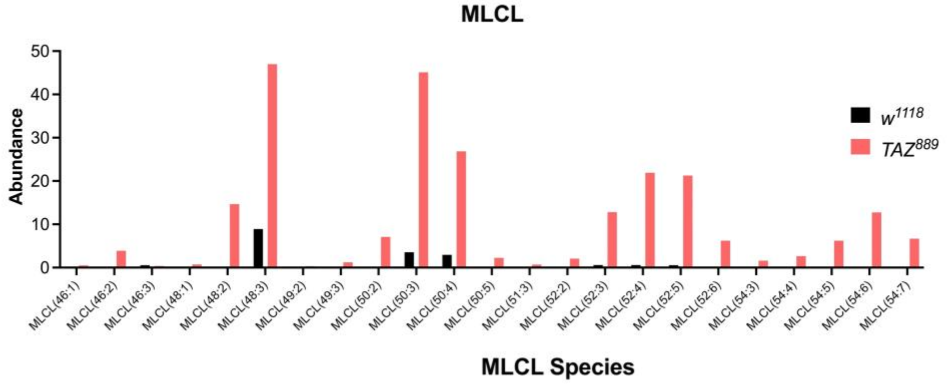


**Supplementary Figure 2: The CL and MLCL species in *w*$^{1118}$ and *TAZ*$^{889}$ flies.** (A.) The most abundant CL species in *w*$^{1118}$ flies are 64:4 and 66:5. The amount of those species is reduced in *TAZ*$^{889}$ flies (Two-tailed Student's *t*-test, n=6 flies per biological replicate, data are mean±s.d.). (B.) Multiple MLCL species accumulate in *TAZ*$^{889}$ flies.

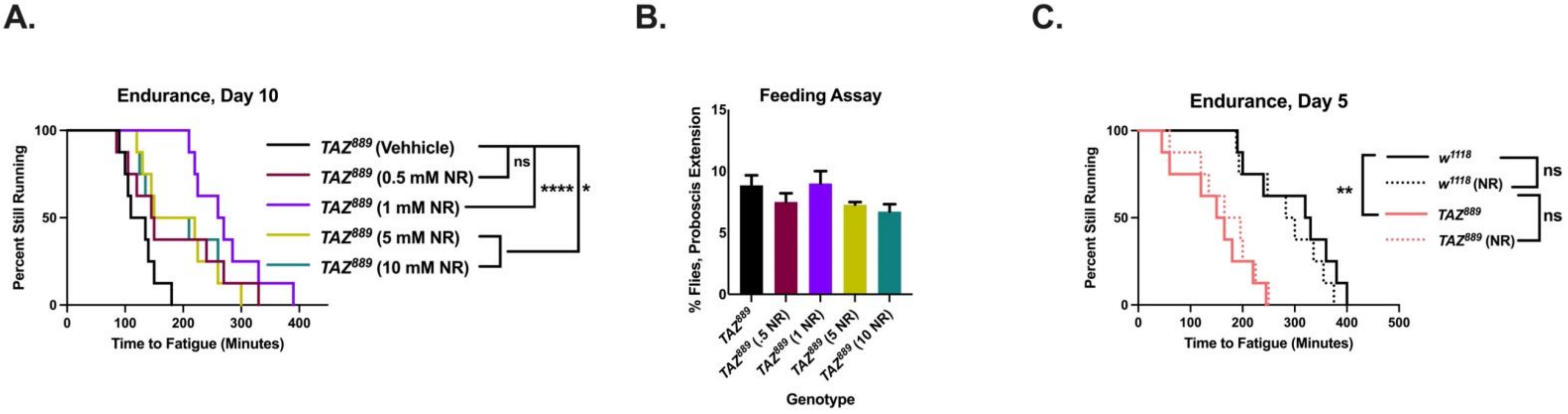


**Supplementary Figure 3: The effects of the timing and dosage of NR supplementation.** (A.) Multiple NR concentrations improved the endurance of *TAZ*$^{889}$ flies, but 1mM provided the biggest benefit (log-rank analysis, n=8 vials, 20 flies per vial). (B.) To rule out differences in feeding behavior contributing to this result, a proboscis extension assessment was performed to measure feeding rate (Wong et al., 2009). There was no significant difference in the percent of flies feeding between different NR concentrations (2 biological replicates ±s.d., n=20 per replicate, One-way ANOVA, p=0.08). (C.) Feeding 1mM of NR for three days does not cause an improvement to the endurance of *TAZ*$^{889}$ flies (log-rank analysis, n=8 vials, 20 flies per vial). (D.) At age day 10, NR does not change the mt DNA copy number in *w*$^{1118}$ flies. (n= 3 biological replicates, data are mean±s.d., Student's *t*- test).*p<.05, **p<.01, ***p<.001, ****p<0.0001, ns= not significant

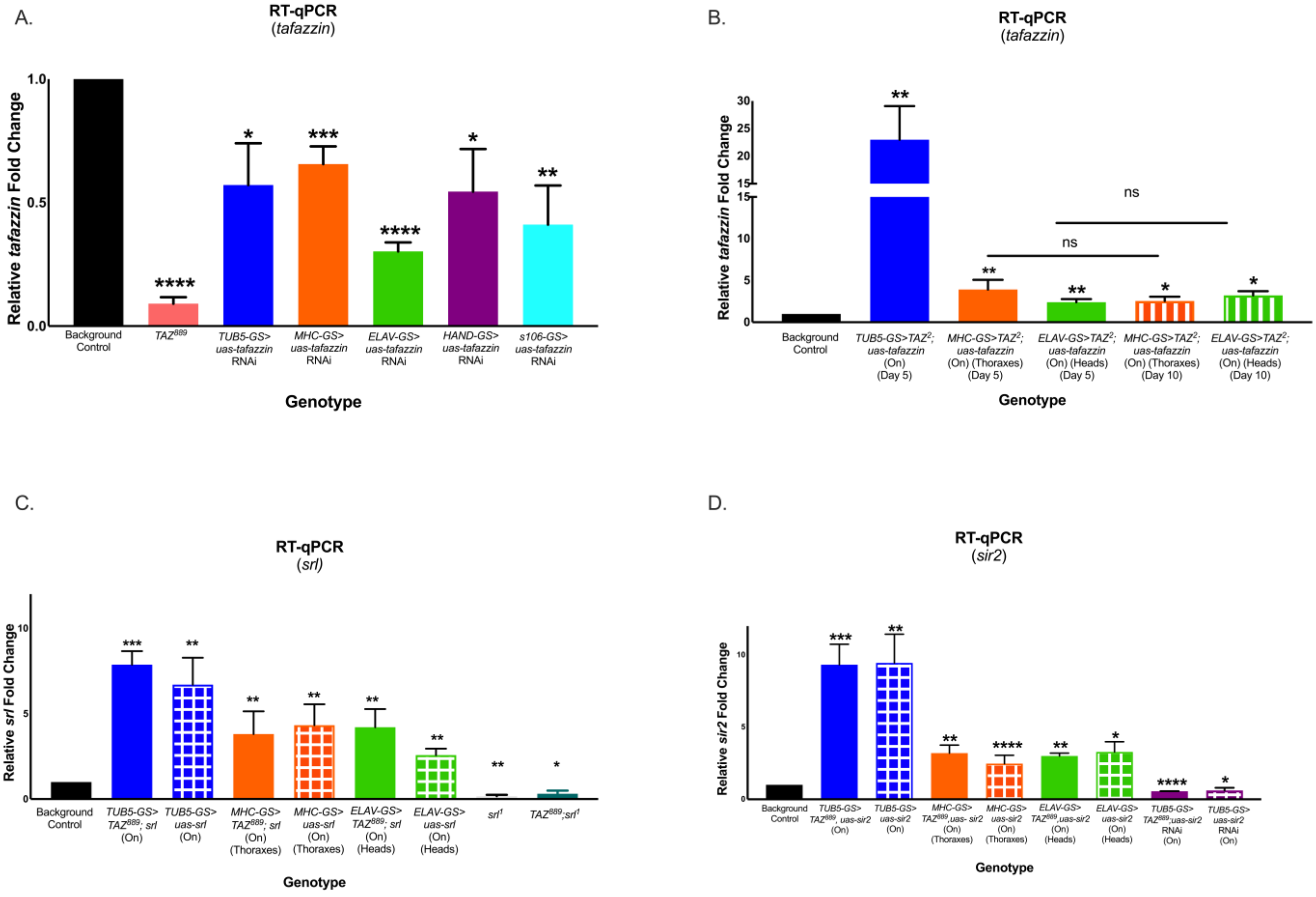


**Supplementary Figure 4: Confirmation of gene expression in target tissues by RT-qPCR.** Bars display the average relative gene expression from triplicate biological samples for (A.,B) *tafazzin*, (C) *srl,* (D), and *sir2.* The "ns" bars show no significant difference in transcript levels between those specific groups. Relative gene expression is expressed in relation to the genetic background control, which in most cases was the uninduced RU- controls. For simplicity, all genetic background control lines' relative gene expression is represented as a value of 1 on the graphs. Relative gene expression was calculated using the ΔΔCT method and analyzed using a two-tailed Student's *t*-test. See methods for more details regarding the isolation procedures, primer sequences and reaction conditions. *p<.05, **p<.01, ***p<.001, ****p<0.0001, ns= not significant, On=mifepristone induced gene expression

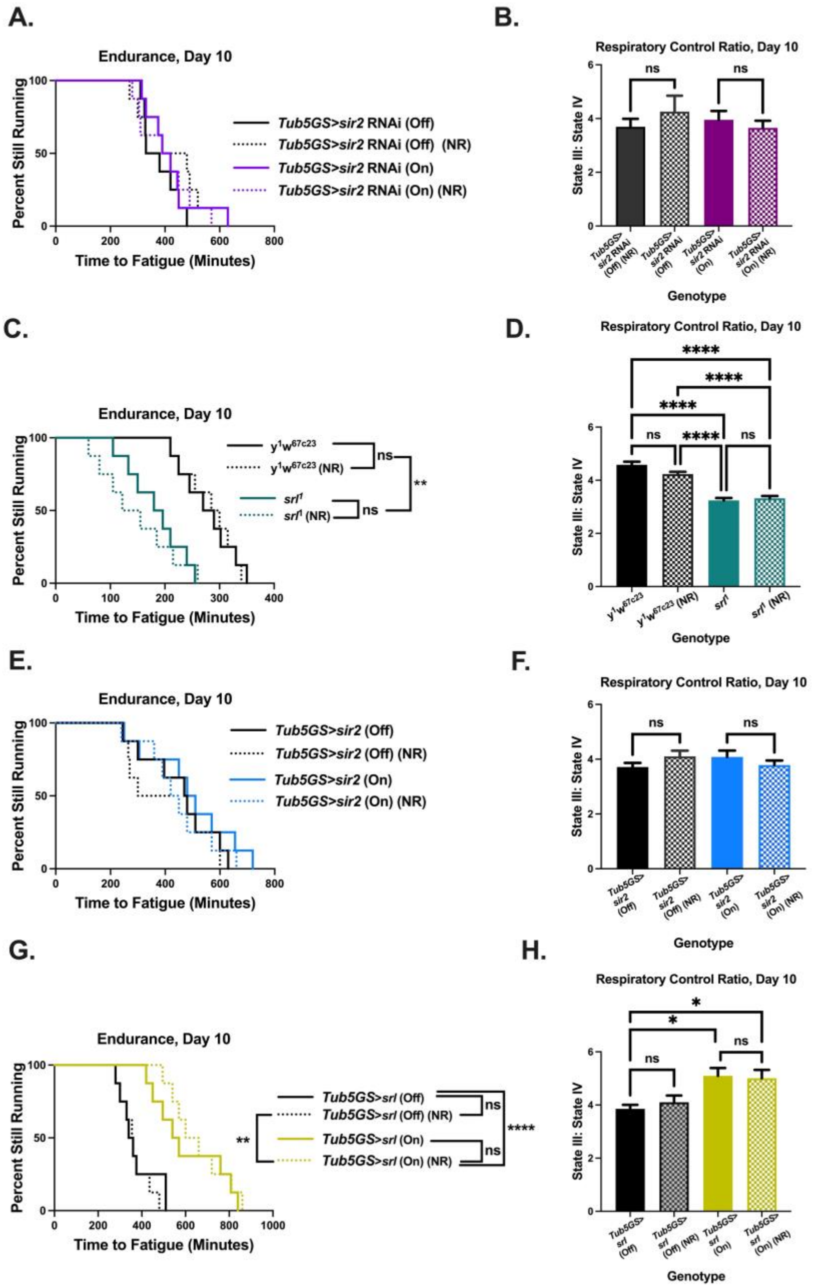


**Supplementary Figure 5: The effects NR and changing *sir2* and *srl* gene expression in control flies.** (A.,B.) Regardless of NR supplementation, knocking down *sir2* does not significantly affect the endurance (n=8 vials, 20 flies per vial) or the RCR (data averages of 6 biological replicates ±s.e.m, n=60 per replicate, 2-way ANOVA, genotype effect, p=0.295, NR effect, p=0.421). (C., D.) *srl*$^{1}$ flies have reduced endurance relative to their background controls (log-rank analysis, p=0.02, n=8 vials, 20 flies per vial), and NR supplementation does not rescue their endurance (data averages of 6 biological replicates ±s.e.m, n=60 per replicate, 2-way ANOVA, genotype effect, p<0.0001, NR effect, p=0.183). (E.,F.) Overexpression of *sir2* has not significant effect on endurance (n=8 vials, 20 flies per vial) or on the RCR (data averages of 6 biological replicates ±s.e.m, n=60 per replicate, 2-way ANOVA, genotype effect, p=0.901, NR effect, p=0.819). (G.,H.) Overexpressing *srl* significantly increase the endurance of control flies (log-rank analysis, p=0.0037, n=8 vials, 20 flies per vial), but NR supplementation does not add to that effect (p=0.456). The RCR is also increased in *srl* overexpressing flies, but there is not additive effect with NR supplementation (data averages of 6 biological replicates ±s.e.m, n=60 per replicate, 2-way ANOVA, genotype effect, p=0.0006, NR effect, p=0.787). p<.05, **p<.01, ***p<.001, ****p<0.0001, ns= not significant, On=mifepristone induced gene expression

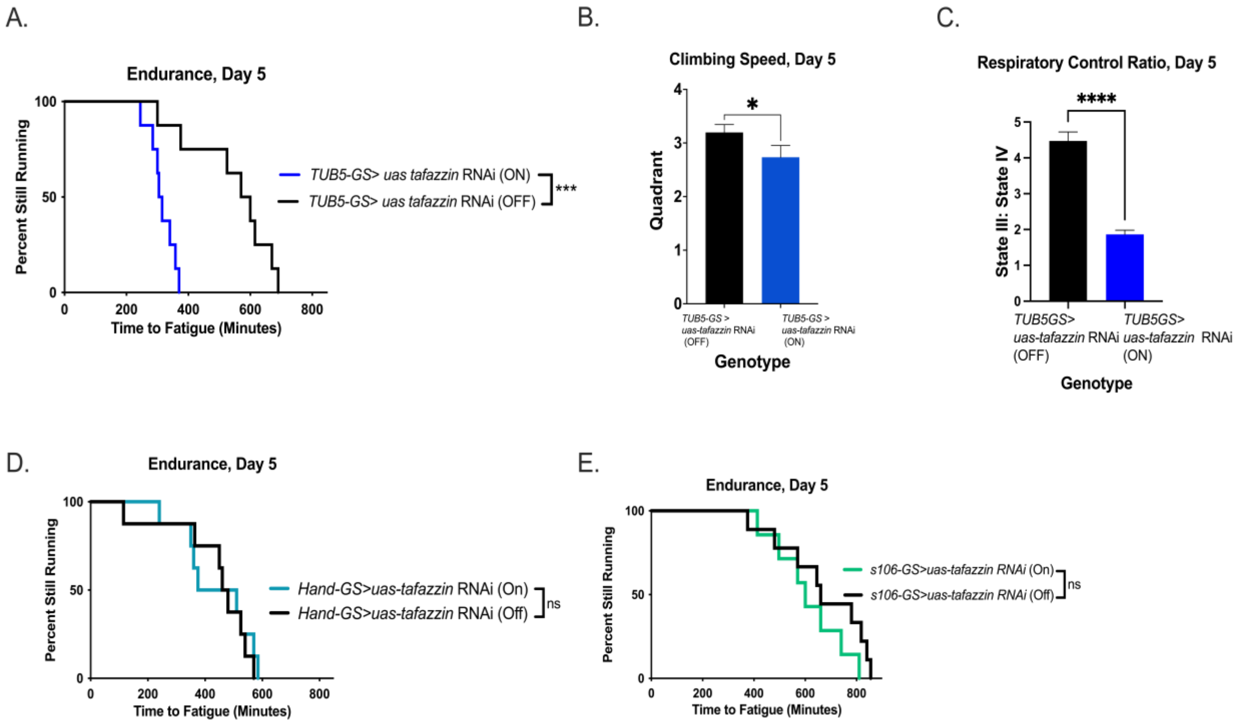


**Supplementary Figure 6: *tafazzin* is not required in heart or fat body tissue of *Drosophila* for normal endurance.** (A.-C.) Ubiquitously knocking down *tafazzin* reduces endurance (log-rank analysis, n=8 vials, 20 flies per vial), slows climbing speed (Two-tailed Student's *t*-test, n= 100 flies, data are mean±s.d.), and lowers the RCR (Two-tailed Student's *t*-test, six biological replicates, n=60 per replicate, data are mean±s.e.m.).(D.) Knocking down *tafazzin* in the heart of adult *Drosophila* does not cause reduced endurance at age Day 5 (log-rank analysis, n=8 vials, 20 flies per vial, p=0.736). (E.) Knocking down *tafazzin* in the fat body of adult *Drosophila* does not lower their endurance at age Day 5 (log-rank analysis, n=8 vials, 20 flies per vial, p=0.2134). *p<.05, **p<.01, ***p<.001, ****p<0.0001, ns= not significant, On=mifepristone induced gene expression

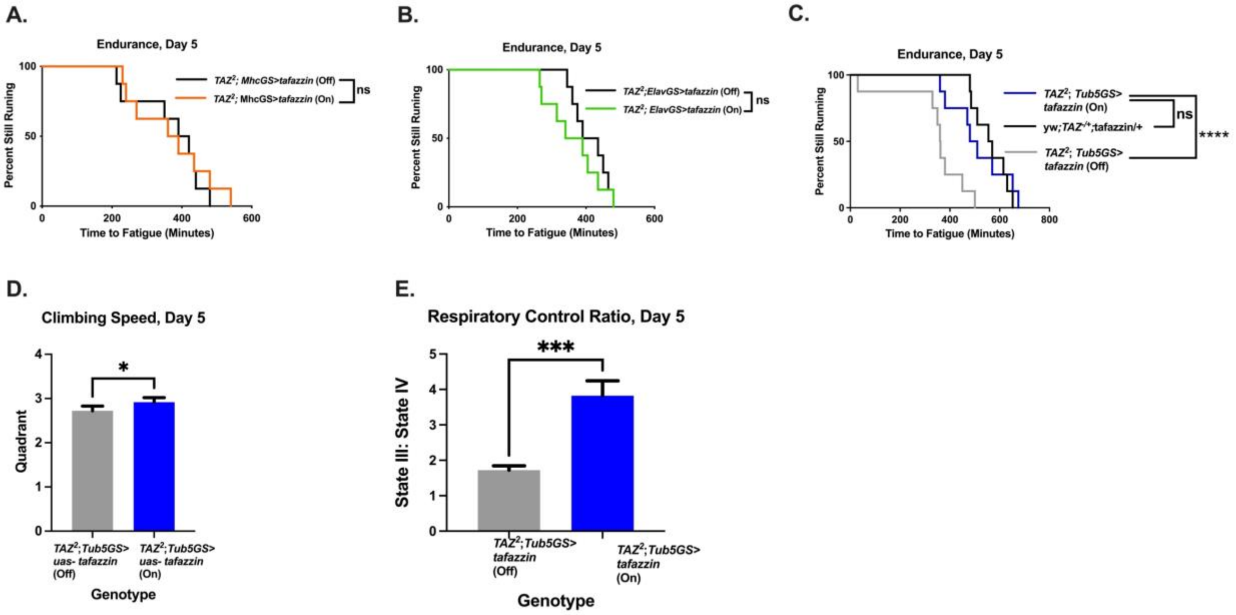


**Supplementary Figure 7: Overexpression of *tafazzin* in the muscle or neuronal tissue of *TAZ* mutants does not rescue endurance at age day 5.** (A.,B.) At age day 5, there is no improvement in the endurance of *TAZ* mutants with overexpression of *tafazzin* the muscle tissue (log-rank analysis, n=8 vials, 20 flies per vial, p=0.679) or in the neuronal tissue at age day 5 endurance at age day 5 (log-rank analysis, n=8 vials, 20 flies per vial, p=0.378). At age day 5, ubiquitously overexpressing *tafazzin* is sufficient to rescue the endurance to control levels (*yw;TAZ*$^{+/-}$*, uas-tafazzin/*+, p=0.908), improve the climbing speed, and fully restore the RCR of *TAZ* mutant flies. *p<.05, **p<.01, ***p<.001, ****p<0.0001, ns=not significant, On=mifepristone induced gene expression
